# Molecular characteristics and laminar distribution of prefrontal neurons projecting to the mesolimbic system

**DOI:** 10.1101/2022.02.17.480862

**Authors:** Ákos Babiczky, Ferenc Mátyás

## Abstract

Prefrontal cortical influence over the mesolimbic system – including the nucleus accumbens (NAc) and the ventral tegmental area (VTA) - is implicated in various cognitive processes and behavioral malfunctions. The functional versatility of this system could be explained by an underlying anatomical complexity; however, the detailed characterization of the medial prefrontal cortical (mPFC) innervation of the NAc and VTA is still lacking. Therefore, combining classical retrograde and conditional viral tracing techniques with multiple fluorescent immunohistochemistry, we sought to deliver a precise, cell- and layer-specific anatomical description of the cortico-mesolimbic pathways. We demonstrated that NAc- (mPFC_NAc_) and VTA-projecting mPFC (mPFC_VTA_) populations show different laminar distribution (layers 2/3-5a and 5b-6, respectively) and express different molecular markers. Specifically, calbindin and Ntsr1 are specific to mPFC_NAc_ neurons, while mPFC_VTA_ neurons express high levels of Ctip2 and FoxP2, indicating that these populations are mostly separated at the cellular level. We directly tested this with double retrograde tracing and Canine adenovirus type 2-mediated viral labelling and found that there is indeed minimal overlap between the two populations. Furthermore, whole-brain analysis revealed that the projection patter of these populations is also different throughout the brain. Taken together, we demonstrated that the NAc and the VTA are innervated by two, mostly non-overlapping mPFC populations with different laminar distribution and molecular profile. These results can contribute to the advancement in our understanding of mesocorticolimbic functions and its disorders in future studies.

## Introduction

The medial prefrontal cortex (mPFC), the nucleus accumbens (NAc) and the ventral tegmental area (VTA) are the three major elements of the mesocorticolimbic system that controls a wide range of behaviors (Tzschentke and Schmidt, 2000; Russo and Nestler, 2013; Riga et al., 2014). mPFC provides the major source of glutamatergic input to the NAc (Brog et al., 1993; Asher and Lodge, 2012; Li et al., 2018) and to the VTA (Geisler and Zahm, 2005; Mahler and Aston-Jones, 2012; Faget et al., 2016). Direct mPFC innervation in the NAc has been implicated in various cognitive processes and malfunctions, such as attention regulation (Christakou et al., 2004), impulse control (Feja and Koch, 2015), addiction (Schmidt et al., 2005; Peters et al., 2008; Seif et al., 2013; Domingo-Rodriguez et al., 2020) and depression (Vialou et al., 2014). mPFC can also bidirectionally modulate neuronal activity in VTA, including NAc- and mPFC-projecting dopaminergic neurons (Gariano and Groves, 1988; Carr and Sesack, 2000; Lodge, 2011). Accordingly, stimulation of excitatory neurons in the mPFC elicits dopamine release in the NAc via the VTA (Taber et al., 1995; Karreman and Moghaddam, 1996) and optogenetic activation of mPFC input in the VTA is reinforcing (Beier et al., 2015; Pan et al., 2021). Although, excitatory neurons in the mPFC are distributed in distinct layers and possess various projection patterns and molecular identity, it is not known how this diversity correlates to the above-mentioned cortical functions.

Several well established classification systems exist, based on anatomical, physiological, molecular and connectivity profile of excitatory neurons in primary motor/sensory neocortical areas (Harris et al., 2014, 2019; Harris and Shepherd, 2015; Baker et al., 2018; Bakken et al., 2021). A widely accepted one divides principal neurons to three major classes according to their laminar distribution and projection pattern. Intertelencephalic (IT) cells are present in layers 2-6 (L2-6) and project to ipsi- and contralateral neocortex and striatum. Neurons of the pyramidal tract (PT, also known as extratelencephalic) class are located mostly in the L5b and innervate mostly mesencephalic and diencephalic regions. The third, corticothalamic (CT) class is composed of neurons in the L6 that innervate the thalamus. However, some studies suggest that this classification might be oversimplified and not universally applicable to all cortical areas (Groh et al., 2010; Kim et al., 2015). Indeed, recent publications divided motor cortical neurons into even more new subclasses according to their genetic identity (Callaway et al., 2021; Zhang et al., 2021). Thus, detailed class-, layer- and cell-selective examination is necessary to validate traditional classification systems of cortical pyramidal neurons, also in the mPFC.

The lack of wide-spread adoption of such specific experimental approaches in the mPFC might be the source of numerous contradictions and inconsistencies present in the mesocorticolimbic literature. For instance, in a recent publication, Kim et al. (2017) demonstrated that NAc- and VTA-projecting mPFC neurons are mostly separated at the cellular level, while other experiments investigating the target selectivity of mPFC cells yielded contradictory results (Thierry et al., 1983; Ferino et al., 1987; Cassell et al., 1989; Pinto and Sesack, 2000; Gabbott et al., 2005; Morishima and Kawaguchi, 2006; Vázquez-Borsetti et al., 2011; Stuber et al., 2011; Kim et al., 2017; Otis et al., 2017; Gao et al., 2020). Such inconsistencies could be resolved by applying combined layer- and cell-selective approaches.

Therefore, combining classical tracing techniques, conditional viral labelling, and multiple fluorescent staining, we have begun to describe the prefrontal innervation of the NAc and VTA in a class-, layer- and cell-specific manner. Here we report that these two structures are innervated by two, rather non-overlapping mPFC neuron populations. While NAc-innervating neurons tend to be found in the L2/3 and L5a, VTA-projecting cells are mostly localized in the L5b and L6, resembling IT and PT projection classes, respectively. Accordingly, these two populations express different combination of molecular markers and have different afferent connections throughout the brain. Furthermore, we found that in comparison with primer cortical areas, the mPFC differs in several cytoarchitectural features.

## Results

### Distribution and molecular characterization of NAc-projecting mPFC cells

In order to investigate the mPFC-NAc connection, first, we injected retrograde tracers Cholera toxin B subunit (CTB) or FluoroGold (FG) into the NAc (Fig. 1*A-C*). Injection sites included both the core (NAcC) and shell (NAcSh) region (Fig. 1*C*). Retrogradely labelled NAc-projecting mPFC cells (mPFC_NAc_) were present throughout the mPFC. To identify the exact subregional distribution of mPFC_NAc_ neurons, we performed multiple fluorescent immunohistochemical (IHC_Fluo_) staining for different molecular markers. As it was previously reported (Mátyás et al., 2014), parvalbumin (PV) staining delineates the dorsal and ventral borders of the prelimbic (PrL) subregion of the mPFC (Fig. 1-1*A*, asterisk). Calbindin (Calb1) was used to define layer 2/3 (L2/3) (van Brederode et al., 1991; Sun et al., 2002) and the ventral border of the infralimbic cortex (IL), where the clearly visible L2/3 diminishes, as well as to visualize the thickening of L1, a characteristic of the deep peduncular cortex (DP) (Akhter et al., 2014) (Fig. 1-1*B*, number sign). COUP-TF-interacting protein 2 (Ctip2, also known as Bcl111b) was used to outline the L5b and L6 (Arlotta et al., 2005; Ueta et al., 2014; Kim et al., 2017) (Fig. 1-1*C*). Furthermore, forkhead box protein P2 (FoxP2) staining identifies the L6 (Ferland et al., 2003) and the gradual thinning and disappearance of a distinct L6 towards the ventralmost part of the mPFC (Fig. 1-1*D*, cross).

**Figure 1.**
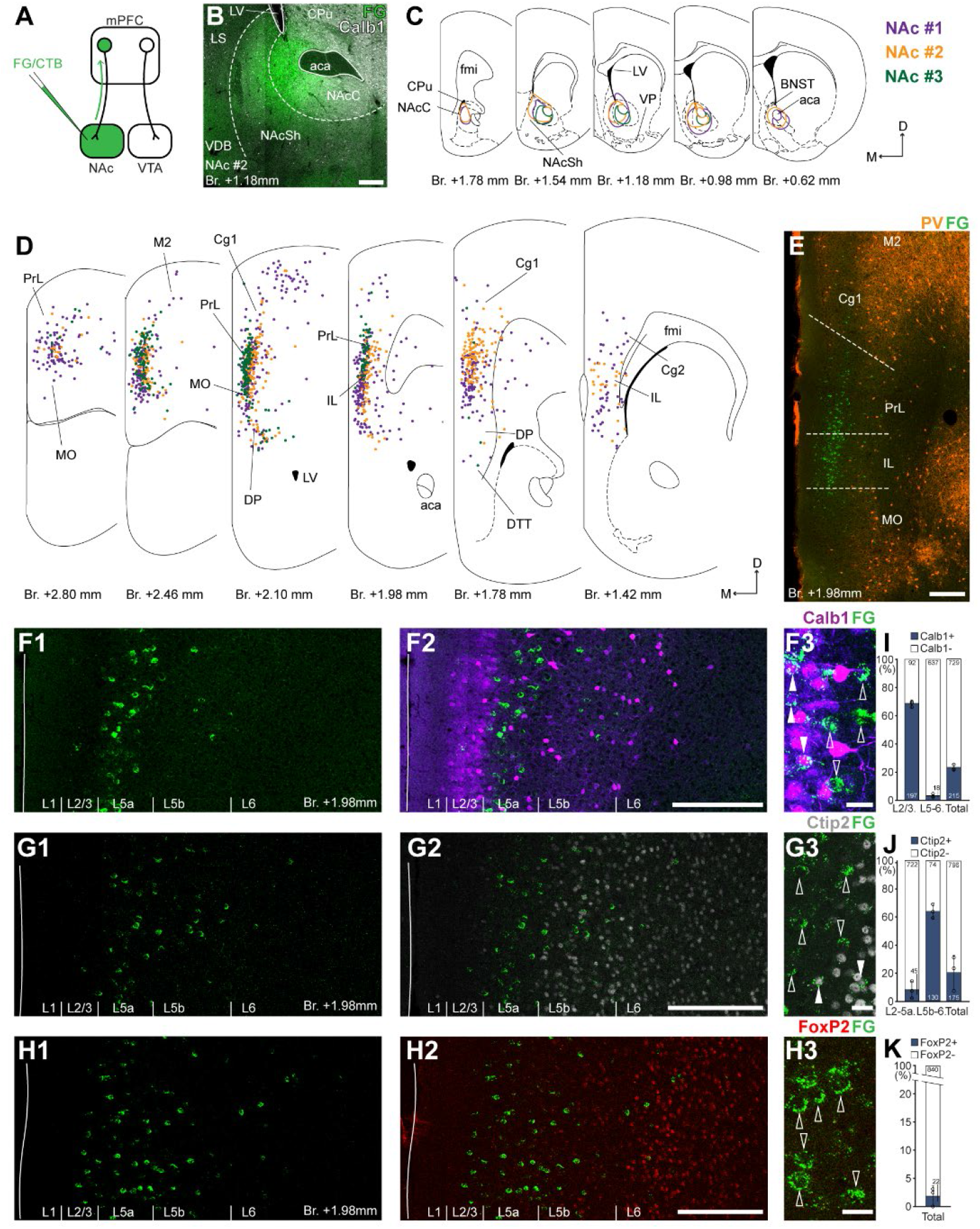
NAc is innervated by L2/3 and L5 mPFC cells. **A,** Experimental design. **B,** A representative retrograde tracer (FG, green) injection site in the NAc. **C,** Delineation of injection sites in the NAc of three animals. Each case is represented with different color. **D,** Plotted distribution of retrogradely labelled cells throughout the mPFC of the same animals as in **C** (same colors represent same animals). Each dot represents one labelled mPFC_NAc_ cell. **E,** Distribution of labelled mPFC_NAc_ neurons in relation to PV (orange) immunofluorescent labelling outlining the PrL cortex (**Fig. 1-1**). **F-H,** Confocal images showing the distribution of FG labelled cells (green) in the PrL (**F1-H1**) with the counterstaining of Calb1 (purple, **F2**), Ctip2 (grayscale, **G2**) and FoxP2 (red, **H2**) (**Fig. 1-1**). Note that most labelled cells are localized in the L2/3 (Calb1) and L5a (Ctip2). **F3-H3,** High magnification confocal images showing the coexpression of FG and Calb1 (**F3**), Ctip2 (**G3**) or FoxP2 (**H3**). White arrowheads indicate coexpression, empty arrowheads indicate the lack of marker expression. **I-K**, Bar graphs showing the proportion of Calb1- (**I**), Ctip2- (**J**) and FoxP2-expressing (**K**) mPFC_NAc_ cells. Numbers in the bars represent cell counts. Scale bars: **B**, **E**, **F1-H1**, **F2-H2**, 200 μm; **F3-H3**, 20 μm. aca, anterior commissure, anterior part; BNST, bed nucleus of the stria terminalis; CPu, caudate putamen; fmi, forceps minor of the corpus callosum; LS, lateral septum; LV, lateral ventricle; VDB, nucleus of the vertical limb of the diagonal band; VP, ventral pallidum. All data are shown as mean ± SD.

According to the obtained molecular-based mPFC map, most mPFC_NAc_ neurons were found in the PrL, medial orbital (MO) and IL subregions and, to a lower extent, in the DP, cingulate area 1-2 (Cg1-2) and secondary motor (M2) cortices. Only a few cells were found in the dorsal tenia tecta (DTT, also known as anterior hippocampal continuation), primary motor (M1) and the adjacent orbital cortices (Fig. 1*D-E*).

Next, we analyzed the laminar distribution of the retrogradely labelled cells using IHC_Fluo_ against Calb1, Ctip2 and FoxP2 (Fig. 1*F-H*, Fig. 1-1). This analysis revealed that most mPFC_NAc_ neurons localized in the Calb1-rich L2/3 and in the L5a, and, to a lower extent, in the L5b (Fig. 1*F-G*). Only a small number of cells were found in the FoxP2-expressing L6 (Fig. 1*H*).

**Figure 1-1.**
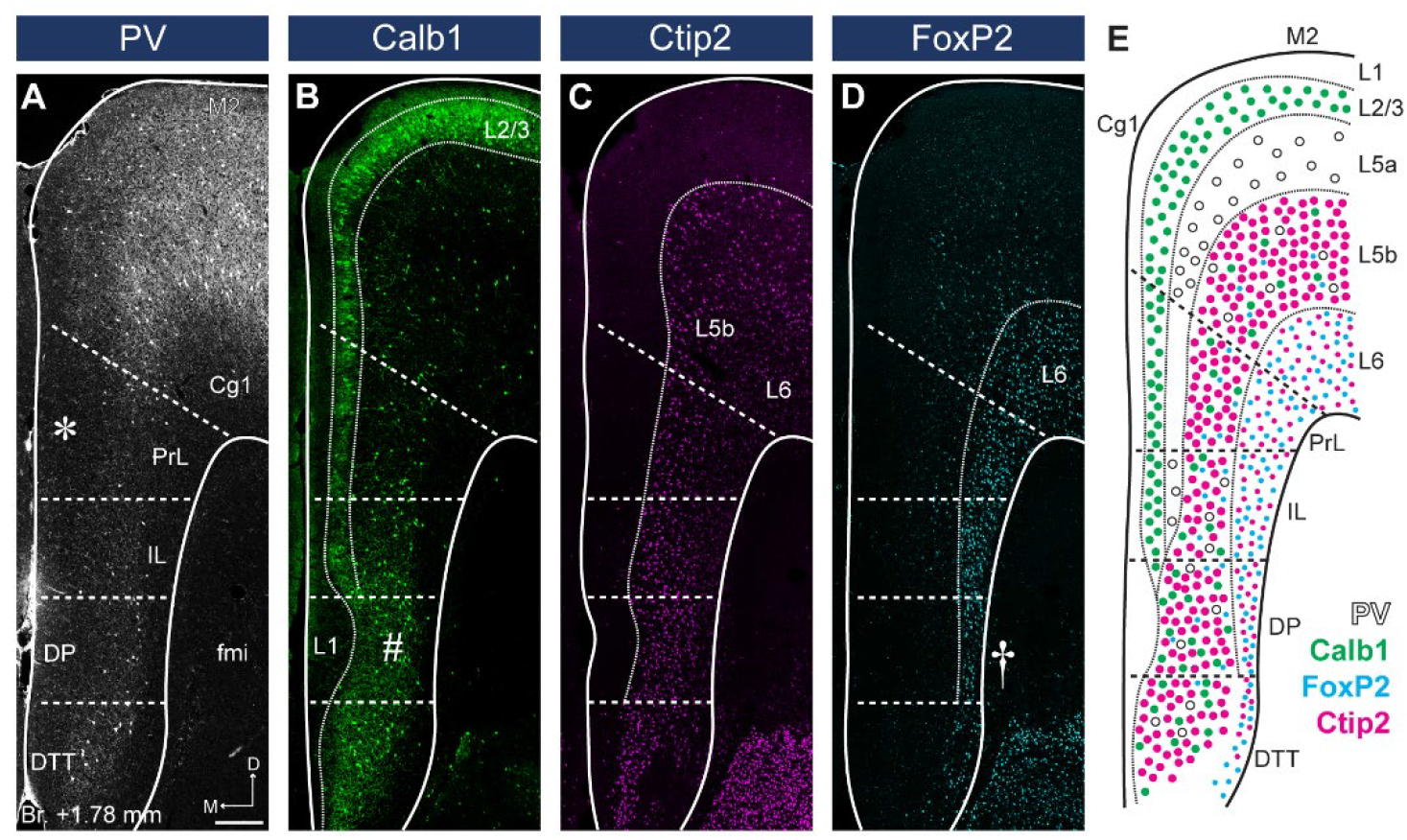
PV, Calb1, Ctip2 and FoxP2 staining define mPFC subregion borders and layers. **A,** The lack of PV+ axonal and cellular immunolabeling (grayscale) in the L2/3 defines the territory of PrL (asterisk). **B,** Calb1-expressing L2/3 (green) is clearly visible in most parts of the mPFC. A thickened L1 and a more compact L2/3 and L5 identifies the DP (number sign). **C,** Ctip2 (magenta) is expressed in the L5b (bigger neurons) and L6 (smaller neurons). **D,** FoxP2 (cyan) is expressed throughout the L6 of the mPFC. Narrowed L6 and the lack of FoxP2-expressing cells (cyan) in the L5 reveal DP (cross). **E,** Schematic summary of PV, Calb1, FoxP2 and Ctip2 distribution in the mPFC. Scale bar: 200 μm.

To characterize the molecular identity of mPFC_NAc_ cells, we quantified their Calb1-, Ctip2 and FoxP2-expression (Fig. 1, *F3-H3, J-K*). Our analysis revealed that about two-thirds (68.64 ± 2.62%, *n* = 3 animals, *N_Calb1+/FG+_* = 197/289 cells; Fig. 1*F3, I*, *left bar*; Table 1) of mPFC_NAc_ neurons in the L2/3 expressed Calb1, while only a small proportion did so in the L5-6 (2.87 ± 1.15%, *N_Calb1+/FG+_* = 18/655 cells; Fig. 1*I*, *middle bar*; Table 1). Collectively, approximately one-fifth of all mPFC_NAc_ neurons expressed Calb1 (22.78 ± 1.86%, *N_Calb1+/FG+_* = 215/944 cells; Fig. 2*I*, *right bar*; Table 1). Although most of the mPFC_NAc_ cells were found in the Ctip2-negative L2/3 and 5a, some cells were found in the deeper layers as well. Confocal analysis revealed that only a small proportion of superficial (i.e., L2/3-5a) cells are Ctip2-positive (8.26 ± 2.6%, *n* = 3 animals, *N_Ctip2+/FG+_* = 45/767 cells; Fig. 1*G3, J*, *left bar*; Table 1), while in the deeper layers (i.e., L5b-6), although relatively few in number, the majority of cells expressed Ctip2 (64.1 ± 4.76%, *N_Ctip2+/FG+_* = 130/204 cells; Fig. 1*J*, *middle bar*; Table 1). Collectively, approximately one-fifth of all mPFC_NAc_ cells expressed Ctip2 (20.8 ± 12.1%, *N_Ctip2+/FG+_* = 175/971 cells; Fig. 1*J*, *left bar*; Table 1). Finally, only a negligible number of mPFC_NAc_ cells expressed FoxP2 (2.11 ± 1.84%, *n* = 3 animals, *N_FoxP2+/FG+_* = 22/862 cells, Fig. 1*H3, K*; Table 1).

**Table 1.** Proportion of FoxP2-, Ctip2- and Calb1-expressing neurons in the mPFC_NAc_ population. * When total cell count was <10, percentage was not calculated. #, number of labelled cells.

| mPFC<sub>NAC</sub> |  | Layers |  |  |  | Total |
| --- | --- | --- | --- | --- | --- | --- |
|  |  | 2/3 | 5a | 5b | 6 |  |
| Calb1 | # FG+ /animal | 128, 101, 60 |  |  | 287, 200, 168 | 415, 301, 228 |
|  | # Calb1+ /animal | 84, 71, 42 |  |  | 7, 4, 7 | 91, 75, 49 |
|  | % Calb1+ (AVG ± SD) | 68.6 ± 2.6% |  |  | 2.9 ± 1.1% | 22.8 ± 1.9% |
| Ctip2 | # FG+ /animal | 62, 356, 349 |  |  | 37, 29, 138 | 99, 385, 487 |
|  | # Ctip2+ /animal | 9, 9, 27 |  |  | 22, 20, 88 | 31, 29, 115 |
|  | % Ctip2+ (AVG ± SD) | 8.26 ± 6.01% |  |  | 64.1 ± 4.76% | 20.8 ± 12.1% |
| FoxP2 | # FG+ /animal | 288, 357, 173 |  | 20, 24, 0 |  | 308, 381, 173 |
|  | # FoxP2+ /animal | 5, 10, 0 |  | 4, 3, 0 |  | 9, 13, 0 |
|  | % FoxP2+ (AVG ± SD) | 1.5 ± 1.4% |  | -* |  | 2.1 ± 1.8% |

Altogether, retrograde tracing experiments revealed that mPFC_NAc_ neurons were mostly localized in the L2/3 and 5a of the PrL, MO and IL cortices. Approximately one-fifth of these cells express Calb1 – most of them are localized in the L2/3, where Calb1 expression is higher (^~^70%) –, and another one-fifth express Ctip2, mostly in the L5b-6.

### Distribution and molecular characterization of VTA-projecting mPFC cells

Next, we investigated the distribution of VTA-projecting neurons in the mPFC (mPFC_VTA_). We used the previously described retrograde tracing approach in the VTA (Fig. 2*A*) identified with IHC_Fluo_ against tyrosine hydroxylase (TH; Fig. 2*B-C*) (Oades and Halliday, 1987; Morales and Margolis, 2017). Most mPFC_VTA_ neurons were localized in the PrL, MO, Cg1-2, IL and DP cortices (Fig. 2*D-E*). There were also several labelled cells in the adjacent orbital and motor cortices as well as in the DTT (Fig. 2*D*).

**Figure 2.**
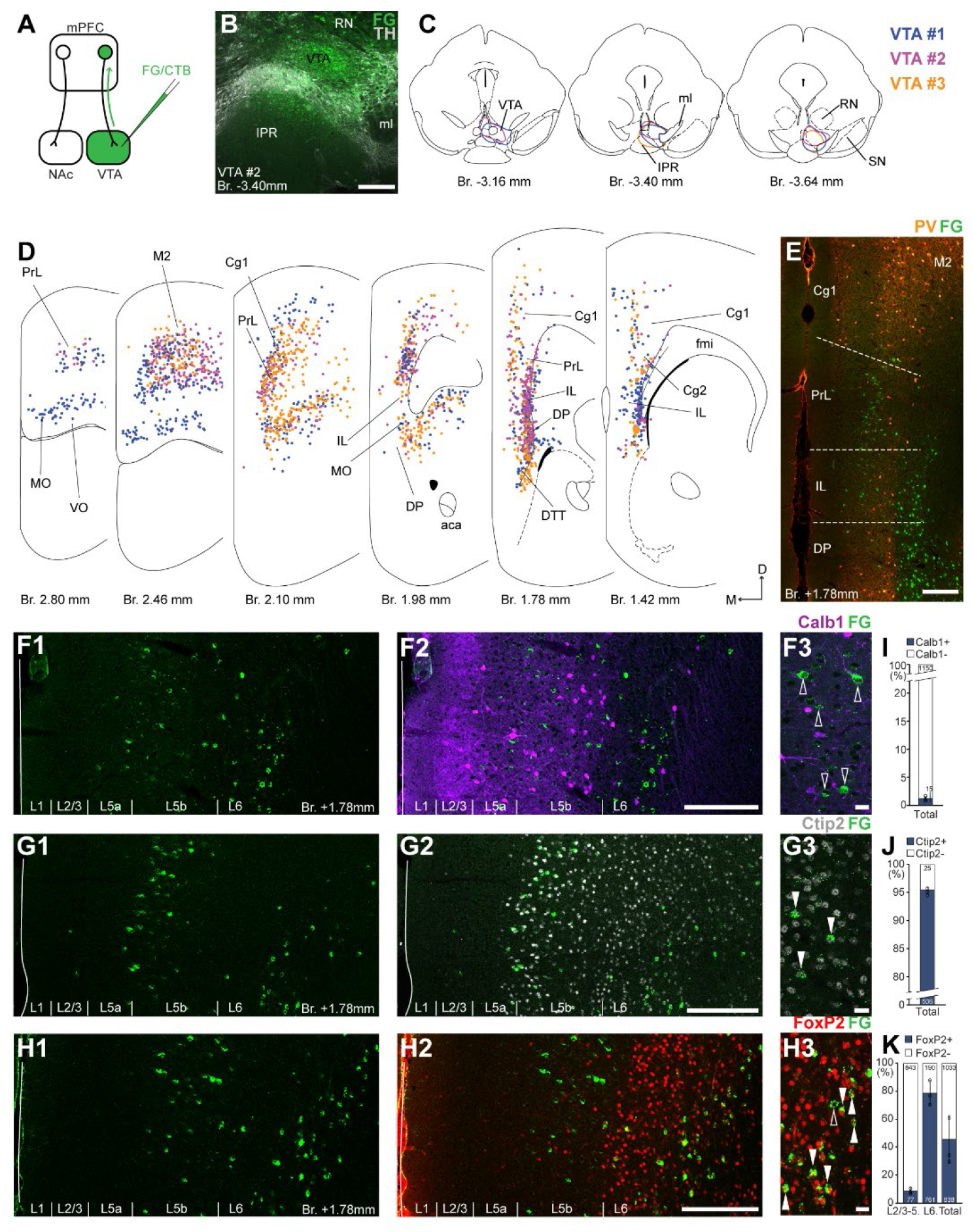
VTA is innervated by two mPFC cell clusters. **A,** Experimental design. **B,** A representative retrograde tracer (FG, green) injection site in the VTA. **C,** Full extent of the injection sites in the VTA in three animals. Each case is represented with different color. **D,** Plotted distribution of retrogradely labelled neurons throughout the mPFC of the same animals as in **C** (same colors represent same animals). Each dot represents one labelled mPFC_VTA_ cell. **E,** Distribution of labelled neurons in the mPFC in relation to PV (orange) immunofluorescent labelling outlining the PrL cortex. **F-H,** Confocal images showing the layer-specific distribution of FG labelled cells (green) in the PrL (**F1-H1**) with counterstaining of Calb1 (purple, **F2**), Ctip2 (grayscale, **G2**) and FoxP2 (red, **H2**). Note that the labelled cells are almost exclusively localized in the L5b (Ctip2) and L6 (Ctip2+FoxP2) layers. **F3-H3,** High magnification confocal images showing the coexpression of FG and Calb1 (**F3**), Ctip2 (**G3**) or FoxP2 (**H3**). White arrowheads indicate coexpression, empty arrowheads indicate the lack of marker expression. **I-K,** Bar graphs showing the proportion of Calb1- (**I**), Ctip2- (**J**) and FoxP2-expressing (**K**) mPFC_VTA_ cells. Numbers in the bars represent cell counts. Scale bars: **B**, **E**, **F1-H1**, **F2-H2**, 200 μm; **F3-H3**, 20 μm. aca, anterior commissure, anterior part; fmi, forceps minor of the corpus callosum; IPR, interpeduncular nucleus, rostral subnucleus; ml, medial lemniscus; RN, red nucleus; SN, substantia nigra; VO, ventral orbital cortex. All data are shown as mean ± SD.

Regarding their laminar distribution, we found that there were two main clusters of mPFC_VTA_ cells (Fig. 2*D, F-H*). One, visually smaller population was found in the more superficial part of the posterior MO, PrL and Cg1, and throughout the Cg2, IL and DP cortices (Fig. 2*D*). Cells in this cluster were mostly localized close to the border of L5a and L5b (Fig. 2*F-G*). The other cluster was found in the deeper parts (L5b-L6) of the anterior MO, PrL and Cg1, and posterior IL and DP cortices (Fig. 2*D*, *H*). The separation of these two mPFC_VTA_ clusters was most prominent between Bregma +1.2-1.8 mm, as it was also shown in previous publications (Geisler and Zahm, 2005; Mahler and Aston-Jones, 2012).

Higher magnification confocal analysis revealed that only a marginal proportion (1.31 ± 0.5%, *n* = 3 animals, *N_Calb1+/FG+_* = 15/1165 cells, Fig. 2*F3, I*; Table 2) of all mPFC_VTA_ cells expressed Calb1. We also quantified the Ctip2-expression of mPFC_VTA_ neurons and found that the vast majority of these cells express Ctip2 (95.07 ± 0.6%, *n* = 3 animals, *N_Ctip2+/FG+_* = 481/506 cells; Fig. 2*G3, J*; Table 2). This finding is in accordance with previous results (Kim et al., 2017) showing CTIP2 gene enrichment in mPFC_VTA_ neurons. Finally, most L6 mPFC_VTA_ cells expressed FoxP2 (78.86 ± 8.79%, *n* = 3 animals, *N_FoxP2+/FG+_* = 761/951 cells; Fig. 2*H3, K*, *middle bar*; Table 2). On the other hand, in the superficial layers (L2/3-L5), only a small proportion (8.69 ± 2.13%, *N_FoxP2+/FG+_* = 77/920 cells; Table 2) of mPFC_VTA_ cells were FoxP2-positive (Fig. 1*K*, *left bar*). In total, about half of all mPFC_VTA_ neurons expressed FoxP2 (45.93 ± 17.15%, *N_FoxP2+/FG+_* = 838/1871 cells; Fig. 2*K*, *right bar*; Table 2).

**Table 2.** Proportion of FoxP2-, Ctip2- and Calb1-expressing neurons in the mPFC_VTA_ population. * When total cell count was <10, percentage was not calculated. #, number of labelled cells.

| mPFC<sub>VTA</sub> |  | Layers |  |  |  | Total |
| --- | --- | --- | --- | --- | --- | --- |
|  |  | 2/3 | 5a | 5b | 6 |  |
| Calb1 | # FG+ /animal | 4, 2, 5 |  |  | 371, 452, 331 | 375, 454, 336 |
|  | # Calb1+ /animal | 4, 2, 2 |  |  | 3, 2, 2 | 7, 4, 4 |
|  | % Calb1+ (AVG ± SD) | -* |  |  | -* | 1.3 ± 0.5% |
| Ctip2 | # FG+ /animal | 1, 2, 1 |  |  | 152, 163, 187 | 153, 165, 188 |
|  | # Ctip2+ /animal | 1, 0, 0 |  |  | 144, 158, 178 | 145, 158, 178 |
|  | % Ctip2+ (AVG ± SD) | -* |  |  | 95.6 ± 1.2% | 95.1 ± 0.6% |
| FoxP2 | # FG+ /animal | 162, 347, 411 |  | 393, 283, 275 |  | 555, 630, 686 |
|  | # FoxP2+ /animal | 17, 22, 38 |  | 347, 219, 195 |  | 364, 241, 233 |
|  | % FoxP2+ (AVG ± SD) | 8.7 ± 2.1% |  | 78.9 ± 8.8% |  | 45.9 ± 17.2% |

Taken together, using retrograde tracing experiments we identified two clusters of mPFC_VTA_ neurons distributed throughout the mPFC: one FoxP2-, and most probably Ctip2-expressing population localized mostly in the L6 (approximately half of all neurons); and one, mostly FoxP2-negative, but Ctip2-positive population in the layer 5b.

### Utility of Cre mouse lines to label mPFC neurons in a layer-selective manner

We found retrogradely labelled mPFC_NAC_ and mPFC_VTA_ neurons in all cellular layers of the mPFC in varying densities. Next, we sought to confirm the laminar organizations of the projecting cells using transgenic mice expressing Cre-recombinase enzyme in a layer-selective manner. We used the following layer-specific Cre-expressing mouse strains: Calb1- (L2/3), Retinol Binding Protein 4- (Rbp4; L5), Neurotensin Receptor 1 (Ntsr1; L6) and FoxP2-Cre (L6) (van Brederode et al., 1991; Hof et al., 1999; Sun et al., 2002; Ferland et al., 2003; Molyneaux et al., 2007; Harris et al., 2014, 2019; Sundberg et al., 2018; Callaway et al., 2021; Matho et al., 2021; Muñoz-Castañeda et al., 2021) in combination with Cre-dependent adeno-associated viral vectors (AAVs) (Fig. 3*A-E*). Furthermore, we used a Thymocyte differentiation antigen 1 (Thy1)-Cre mouse line as control, in which Cre enzyme is expressed in all pyramidal neurons, regardless of their laminar localization (Fig. 3*F*).

**Figure 3.**
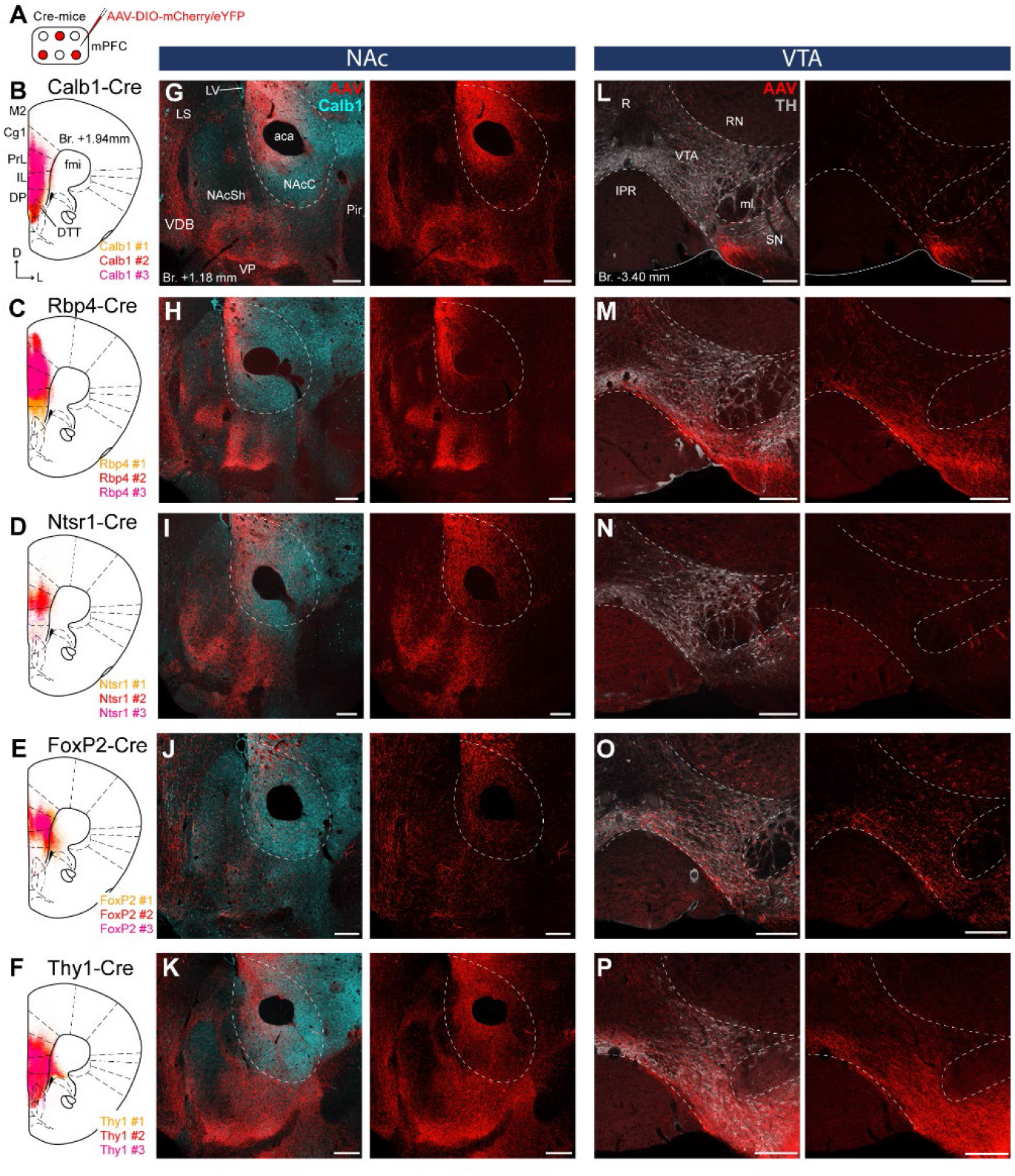
Distinct NAc and VTA innervation by genetically identified mPFC cell populations. **A,** Experimental design. **B-F,** Delineation of viral injection sites in the mPFC of 3-3 animals of the Calb1- (**B**), Rbp4- (**C**), Ntsr1- (**D**), FoxP2- (**E**) and Thy1-Cre (**F**) strains. Viral labelling was always analyzed after immunohistochemical enhancement (**Fig. 3-1**). For detailed distribution of labelled cells in the mPFC and M1 see **Fig. 3-2**. **G-K,** Confocal images showing virally labelled prefrontal axons (red) in the NAc of Calb1-Cre **(G)**, Rbp4-Cre **(H)**, Ntsr1-Cre **(I)**, FoxP2-Cre **(J)** and Thy1-Cre **(K)** mouse strains. Calb1 (cyan) immunofluorescent staining was used to identify the NAcC. **L-P,** Distribution of labelled axons (red) from the same animals (respectively) in the VTA defined with TH staining (grayscale). Scale bars: 200 μm. aca, anterior commissure, anterior part; fmi, forceps minor of the corpus callosum; IPR, interpeduncular nucleus, rostral subnucleus; LS, lateral septum; LV, lateral ventricle; ml, medial lemniscus; Pir, piriform cortex; R, raphe; RN, red nucleus; SN, substantia nigra; VDB, nucleus of the vertical limb of the diagonal band; VP, ventral pallidum.

Virally labelled cell bodies in all strains were primarily found in the PrL, IL, Cg1-2, MO, and, to a lower extent, in the DP, the ventromedial M2, the dorsal part of the DTT and the medial part of the VO cortex (Fig. 3*B-F*) in good correspondence with the distribution of the retrogradely labelled mPFC_NAC_ and mPFC_VTA_ neurons (Fig. 1, Fig. 2). Note that viral expression was always analyzed after IHC-enhancement of eYFP/mCherry, because this method revealed structures – mostly thin axon-branches, but also some cell bodies - and fine details (e.g., dendritic spines) otherwise not detectable (see Methods) (Fig. 3-1).

**Figure 3-1.**
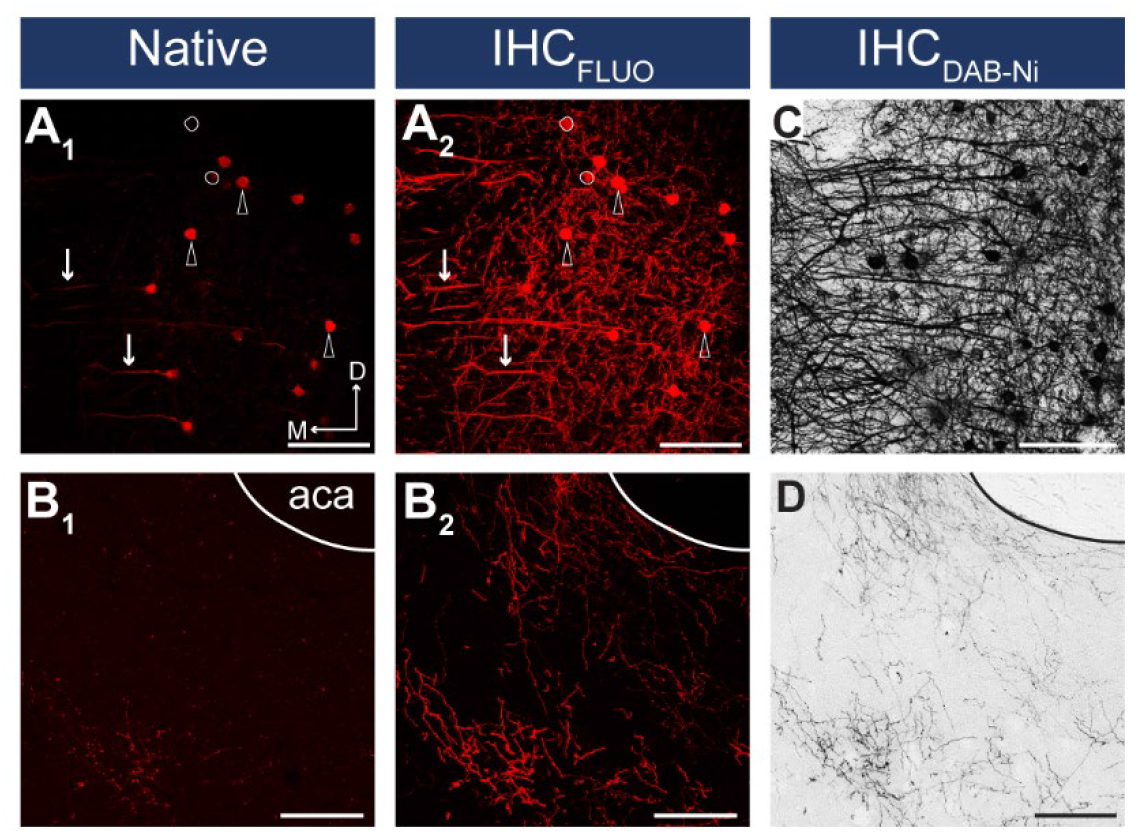
IHC enhancement is necessary for reliable detection of viral fluorescent signal. **A-B,** Confocal images showing the native mCherry signals in the somatodendritic (**A1**) and axonal compartments (**B1**) and after IHC_Fluo_ signal-enhancement (**A2-B2**) of the same slice taken from the mPFC (**A**, injection site) and from the NAc (**B**). Arrows indicate dendrites, arrowheads indicate neurons visible in both the native and IHC_Fluo_-enhanced sample. White circles indicate neurons not visible in the native sample. **C-D,** Brightfield images showing IHC_DAB-Ni_-enhanced samples taken from a neighboring slice. Scale bars: 100 μm. aca, anterior commissure, anterior part.

Since the majority of previous publications describing cortical layer-specific markers focused on primary cortical areas, we compared the expression pattern of virally labelled cells in the mPFC – a higher-order cortical region - (Fig. 3-2*A-E*) and in the primary motor cortex (M1, Fig. 3-2*F-J*) - a primary frontal cortical area – in each mouse strain. Labelled cells in the Calb1-Cre animals showed similar distribution in both cortical areas: most of them were found in the L2/3 with scattered cells in the L5 (Fig. 3-2*A, F*) (Muñoz-Castañeda et al., 2021). Interestingly, Rbp4-, and Ntsr1-expressing cells showed somewhat different distribution in the two cortical regions (Fig. 3-2*B-C, G-H*). In the Rbp4-Cre strain, virally labelled cells in the mPFC were found to some extent in the L2/3 – especially in the ventral part of the mPFC, in the IL and DP – besides the well-known L5 location. In the M1, only the L5 population was present (Callaway et al., 2021; Muñoz-Castañeda et al., 2021) (Fig. 3-2*B, G*). In the Ntsr1-Cre animals, no virally labelled neurons were found in the L6 in the mPFC, only in the L5a (Fig. 3-2*C*). In the M1 cortex, Ntsr1-expressing labelled cells were found exclusively in the L6, as it was previously reported (DeNardo et al., 2015; Tasic et al., 2016; Sundberg et al., 2018; Callaway et al., 2021; Muñoz-Castañeda et al., 2021) (Fig. 3-2*H*). In the Foxp2- and Thy1-Cre animals we did not observe any difference between the two cortical regions: AAV transduced cells were found in the L6 (Fig. 3-2*D, I*), or in all cellular layers of the mPFC (Fig. 3-2*E*) and the M1 (Fig. 3-2*J*), respectively.

**Figure 3-2.**
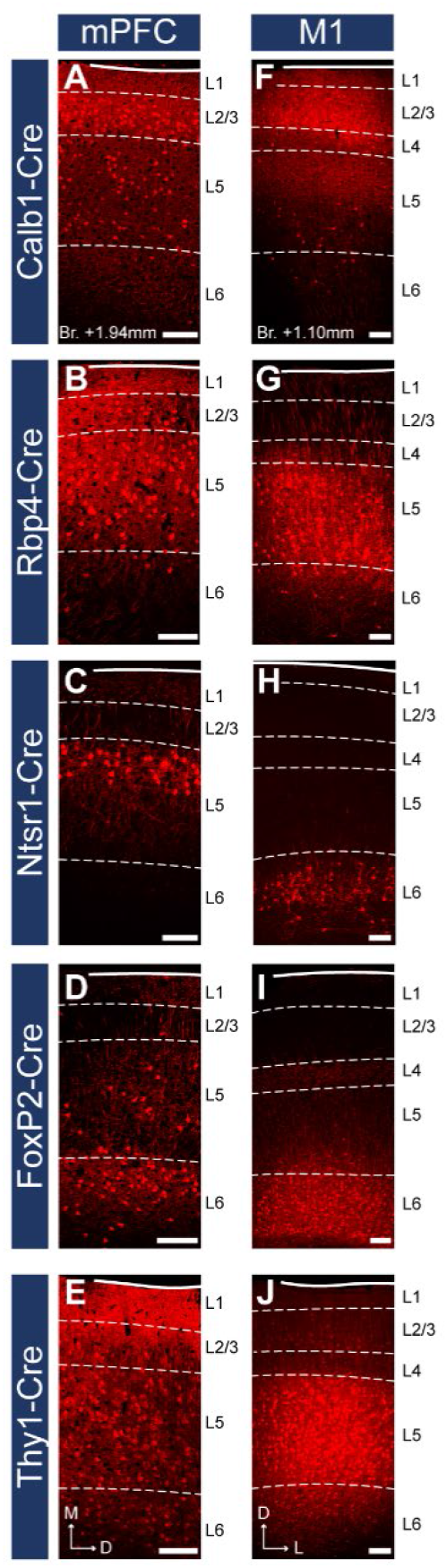
Layer-specific Cre mouse lines reveal different laminal distribution of mPFC and primary motor cortical cells. **A-E,** Confocal images showing the AAV labelled cells in the mPFC in the same animals as in **Fig. 3** (respectively). **F-J,** Localization of labelled neurons in the M1 cortex in the same mouse strains. Note the different distribution of Rbp4- (**B, G**) and Ntsr1-expressing (**C, H**) cells in the two cortical regions. Dashed lines indicate layer borders defined with IHC_Fluo_ against Calb1, Ctip2 or FoxP2 (**Fig. 1-2**). Scale bars: 200 μm.

Together, these results show that these mouse strains can be used to label and investigate distinct layers of prefrontal cell populations, confirming previous findings (van Brederode et al., 1991; Hof et al., 1999; Sun et al., 2002; Ferland et al., 2003; Molyneaux et al., 2007; Harris et al., 2014, 2019; Sundberg et al., 2018; Callaway et al., 2021; Matho et al., 2021; Muñoz-Castañeda et al., 2021). However, in some cases (Rbp4-, Ntsr1-Cre) the distribution of labelled neurons was somewhat different in the mPFC compared to M1.

### Layer-selective prefrontal cortical innervation of the NAc and VTA

After validating the use of these Cre mouse strains and AAV vectors to label mPFC neuron populations in a layer-selective manner, we sought to explore their projection patterns in the NAc and VTA. In order to do this, we performed confocal microscopy combined with multiple IHC_Fluo_ in tissue samples taken from the mPFC animals described in the previous section.

In the Calb1-Cre strain – where viral transduced cells were confined to the L2/3 – labelled axons were found in the NAc (Fig. 3*G*) but not in the VTA (Fig. 3*L*). These results are in accordance with our retrograde tracing results showing that a high proportion of mPFC_NAC_ neurons in the L2/3 express Calb1 (Fig. 1), and the lack of mPFC_VTA_ cells in the Calb1-rich layer 2/3 (Fig. 2). In the Rbp4-Cre animals (L2/3-5), AAV labelled axons were found both in the NAc (Fig. 3*H*) and VTA (Fig. 3*M*) also confirming our retrograde tracing result (Fig. 1, Fig. 2). Ntsr1-Cre expressing cells – localized in the L5a (Fig. 3-1*C*), where most of the mPFC_NAc_ neurons were found previously (Fig. 1) – projected to the NAc with visually dense arborization (Fig. 3*I*) but avoided the VTA (Fig. 3*N*). In the FoxP2-Cre strain (L6), only a small number of AAV-labeled axons was present in the NAc (Fig. 3*J*), while a relatively dense arborization of labelled axons was found in the VTA (Fig. 3*O*). This is in good accordance with our previous findings demonstrating that only a marginal proportion of mPFC_NAC_ neurons express FoxP2 (Fig. 1), while almost half of all mPFC_VTA_ cells does so (Fig. 2). Finally, in the control Thy1-Cre strain we observed dense axonal arborization both in the NAc (Fig. 3*K*) and VTA (Fig. 3*P*).

Taken together, our classical retrograde and cell type-specific anterograde viral tracing experiments revealed that mPFC_NAc_ and mPFC_VTA_ neuron populations are mostly separated in the L2/3-5a and L5b-6 (respectively), although this separation is not exclusive. Conversely, these populations seem to overlap in the L5, but it is not clear whether a single mPFC neuron projects to both targets simultaneously or shows target selectivity.

### NAc- and VTA-projecting mPFC populations are mostly non-overlapping

Next, we aimed to answer the open question whether a single mPFC can innervate the NAc and VTA simultaneously or not. Although previous studies investigated the target selectivity of mPFC neurons extensively, some reported relatively high ratio of multiple-projection (Thierry et al., 1983; Ferino et al., 1987; Cassell et al., 1989; Vázquez-Borsetti et al., 2011; Rojas-Piloni et al., 2017), while others found the opposite (Pinto and Sesack, 2000; Gabbott et al., 2005; Morishima and Kawaguchi, 2006; Kim et al., 2017; Otis et al., 2017; Cruz et al., 2021). Therefore, we carried out two independent experimental approaches to clarify this issue (Fig. 4).

**Figure 4.**
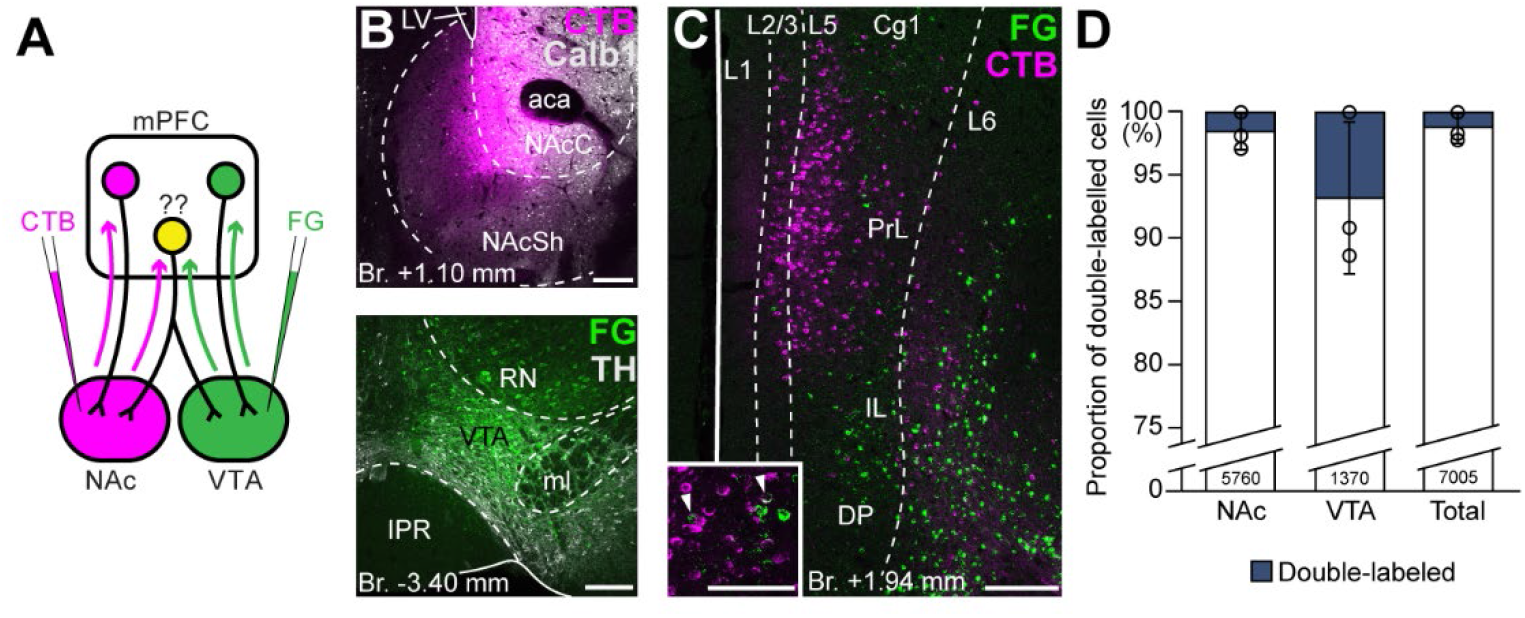
VTA and NAc are mostly innervated by non-overlapping mPFC cell populations. **A,** Experimental design of double retrograde tracing experiments. **B,** Representative CTB (magenta) injection site in the NAc (top) and FG (green) in the VTA (bottom). **C,** High magnification confocal image showing the distribution of mPFC_NAc_ (magenta) and mPFC_VTA_ cells (green) in the mPFC. Inset shows higher magnification of the same slice with arrowheads representing double-labelled cells. **D,** Only a small proportion of labelled mPFC cells innervated both VTA and NAc. Scale bars: 200 μm, **C** inset: 100 μm. aca, anterior commissure, anterior part; IPR, interpeduncular nucleus, rostral subnucleus; LV, lateral ventricle; ml, medial lemniscus; RN, red nucleus. Data are shown as mean ± SD.

First, we performed double retrograde tracings with FG and CTB (interchangeably) from the NAc and the VTA (Fig. 4*A-B*) and investigated the overlap of the labelled populations in the mPFC (Fig. 4*C*). Our results showed that only a small proportion of all cells contained both tracers (NAc+VTA/VTA = 6.78 ± 5.97%, NAc+VTA/NAc = 1.54 ± 1.40%, NAc+VTA/total = 1.26 ± 1.12%; *N_NAc_* = 269, 2940, 2551 cells, *N_VTA_* = 111, 489, 770 cells, *N_VTA+NAc_*= 0, 55, 70 cells; *n* = 3 mice; Fig. 4*D*; Table 3) and most of them were found in the L5.

**Table 3.** Quantification of double retrograde tracing experiments. * Including double-labelled cells. #, number of labelled cells.

|  | NAc* | VTA* | Total* | Double-labelled |
| --- | --- | --- | --- | --- |
| # /animal | 269, 2995, 2621 | 111, 544, 840 | 380, 3374, 3251 | 0, 55, 70 |
| % double-labelled | 1.5 ± 1.4% | 6.8 ± 6.0% | 1.3 ± 1.1% | - |

Although these results indicate that mPFC_NAc_ and mPFC_VTA_ populations are mostly non-overlapping at the cellular level, we considered that double retrograde technique tends to underestimate the actual proportion of multiple projecting cells. Therefore, we also applied an intersectional viral tracing approach to clarify the target selectivity of mPFC neurons. We injected Canine adenovirus type 2 carrying Cre-recombinase gene (CAV2-Cre) into the NAc or VTA, and Cre-dependent AAV-DIO-mCherry into the mPFC (Fig. 5*A-C*), a technique that was previously shown to be suitable to label cortico-tegmental and cortico-accumbal pathways (Beier et al., 2015; Kerstetter et al., 2016; Kim et al., 2017; Cruz et al., 2021). Using this method, we could selectively label mPFC_NAc_ and mPFC_VTA_ neurons with their entire axonal arborization, including collaterals projecting to other brain regions. After confirming that the injection sites were correctly positioned in the NAc or VTA (se*e Methods*, Fig. 5*B1*, *C1*) and in the mPFC (Fig. 5*B2, C2*), we compared the projection pattern of mPFC_NAc_ and mPFC_VTA_ neurons both in the NAc and the VTA (Fig. 5*D-E*). We found that mPFC_NAc_ axons were abundant in the NAc (Fig. 5*D*, *left*), while only a few labelled axons were present in the VTA (Fig. 5*D*, *right*). Conversely, mPFC_VTA_ neurons sent only sparse innervation to the NAc (Fig. 5*E*, *left*), but we found dense innervation in the VTA (Fig. 5*E*, *right*).

**Figure 5.**
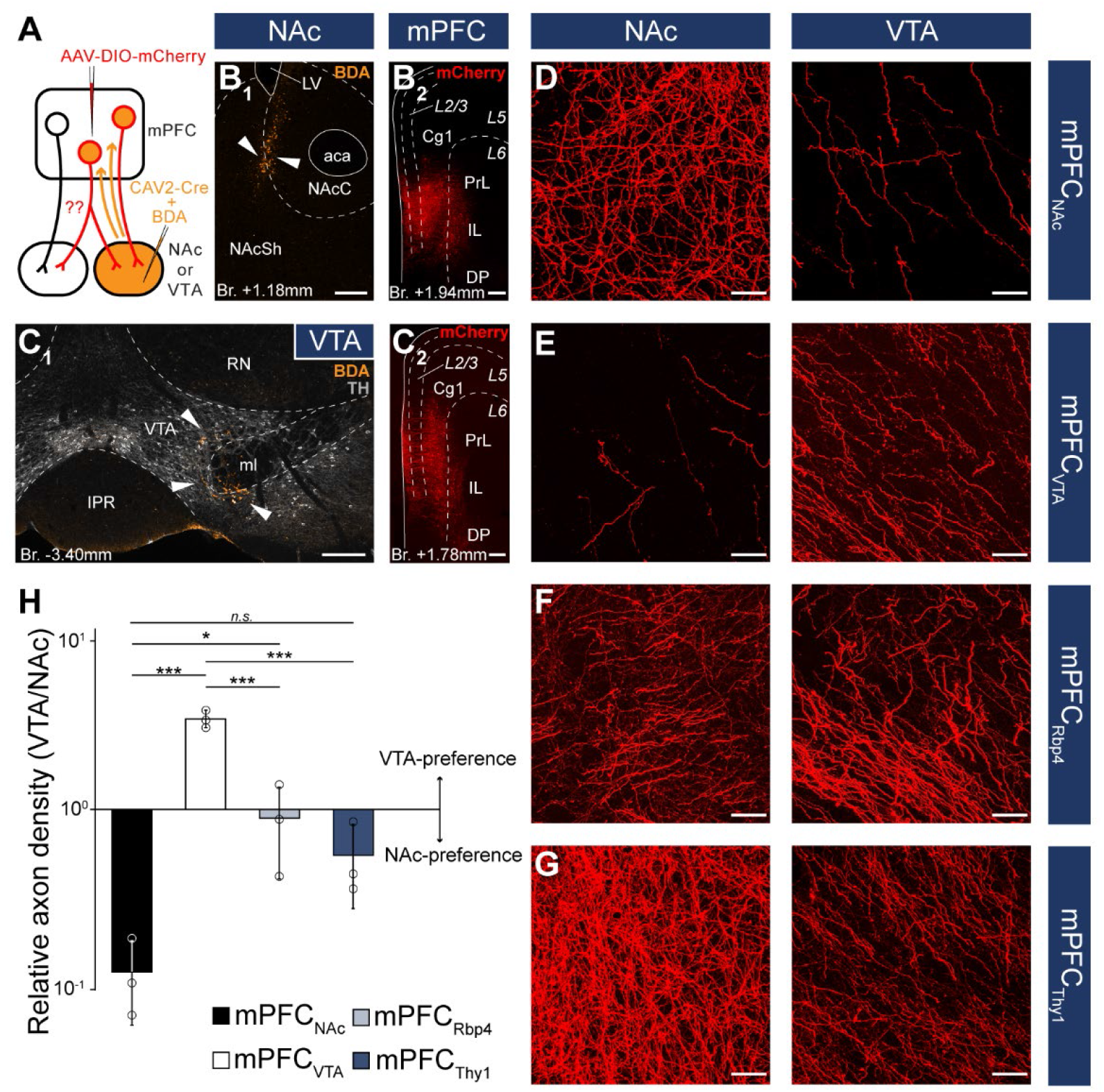
NAc- or VTA-preference of mPFC cells. **A,** Experimental design of CAV2-Cre mediated viral tracing experiments. BDA was used to visualize the exact location of injection sites. **B,** Representative CAV2-Cre + 5% BDA (orange) injection site (**B1**) in the NAc and AAV-DIO-mCherry (red) injection site (**B2**) in the mPFC of the same animal. **C**, Representative CAV2-Cre+BDA (green) injection site (**C1**) in the VTA counterstained with TH (grayscale) and AAV-DIO-mCherry (red) injection site (**C2**) in the mPFC of the same animal. **D-G**, High magnification confocal images showing the distribution of mCherry (red) labelled axons in the NAc (left) and the VTA (right) in a mPFC_NAC_ (**D**), mPFC_VTA_ (**E**), mPFC_Rbp4_ (**F**; same animal as in Fig. 3C, H, M and Fig. 3-1B) and mPFC_Thy1_ (**G**; same animal as in Fig. 3F, K, P and Fig. 3-1E) animal. **H,** Quantification of relative axon density (RAD) in the mPFC_NAC_, mPFC_VTA_, mPFC_Rbp4_ and mPFC_Thy1_ animals. *p < 0.05; ***p < 0.001. Scale bars: **B-C**, 200 μm, **D-G,** 20 μm. aca, anterior commissure, anterior part; BDA, biotinylated dextran amine; IPR, interpeduncular nucleus, rostral subnucleus; LV, lateral ventricle; ml, medial lemniscus; RN, red nucleus. Data are shown as mean ± SD.

To quantify these results, we applied high magnification confocal imaging (63x) to measure and compare the relative axon densities (RAD) in the two target areas. This quantitative analysis showed that mPFC_NAc_ neurons innervated the NAc almost ten-fold stronger than the VTA (RAD_(VTA/NAc)_ = 0.11 ± 0.06; *n* = 3 animals; Fig. 5*H*; Table 4). On the other hand, mPFC_VTA_ cells innervated preferentially the VTA as opposed to the NAc (RAD_(VTA/NAc)_ = 3.45 ± 0.41; *n* = 3 animals; Fig. 5*H*; Table 4).

**Table 4.**
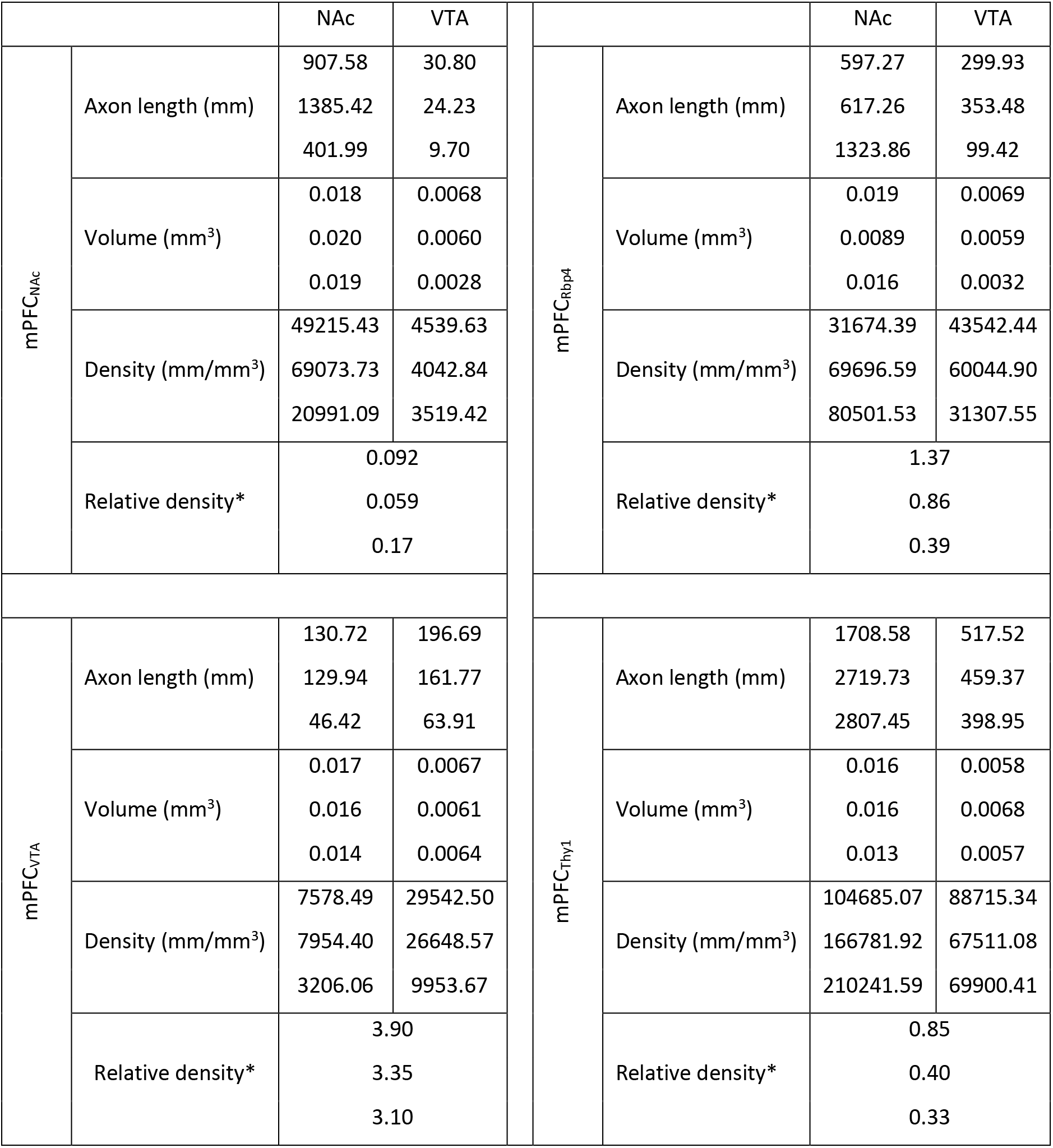
Quantification of axon-length and density in the VTA and NAc in the mPFC_NAc_, mPFC_VTA_, mPFC_Rbp4_ and mPFC_Thy1_ animals. * Relative density = RAD_VTA_/RAD_NAc_.

As controls, we used Rbp4- (mPFC_Rbp4_) and Thy1-Cre (mPFC_Thy1_) animals from the previous viral tracing experiments (Fig. 4), since these two cell populations innervated both the NAc and VTA intensively (Fig. 5*F-G*). Our analysis revealed that mPFC_Rbp4_ cells innervated both regions similarly (RAD_(VTA/NAc)_ = 0.88 ± 0.49; *n* = 3 animals; Fig. 5*F, H*; Table 4), while mPFC_Thy1_ cells tended to innervate NAc slightly more intensively (RAD_(VTA/NAc)_ = 0.53 ± 0.28; *n* = 3 animals; Fig. 5*G-H*; Table 4). One-way ANOVA analysis revealed significant differences between groups (F_(3, 8)_ = 55.56; *p* = 0.000011; mPFC_NAc_ vs mPFC_VTA_, *p* = 0.0000026; mPFC_NAc_ vs mPFC_Rbp4_, *p* = 0.028; mPFC_NAc_ vs mPFC_Thy1_, *p* = 0.18; mPFC_VTA_ vs mPFC_Rbp4_, *p* = 0.000018; mPFC_VTA_ vs mPFC_Thy1_, *p* = 0.0000072; LSD post-hoc test; Fig. 5*H*). These and the double retrograde tracing results indicate that mPFC_NAc_ and mPFC_VTA_ neurons are rather non-overlapping, although there is a marginal population – in the L5 – that innervates both areas.

### mPFC_NAc_ and mPFC_VTA_ populations have different efferent connections

After confirming that mPFC_NAc_ and mPFC_VTA_ neurons are mostly separated at the cellular level, we sought to investigate the projection pattern of these populations throughout the brain. Therefore, we used immunoperoxidase development with DAB-Ni as a chromogen (IHC_DAB-Ni_) (Fig. 3-1) for the mPFC_NAc_ (*n* = 3 mice) and mPFC_VTA_ (*n* = 3 mice) brain samples. Semi-quantitative investigation of the samples revealed clear differences between the two populations (Fig. 6; Table 5). Most notably, mPFC_NAc_ neurons projected intensively to the ipsi- and contralateral striatum – including the NAc (Fig. 6*C*, *left*) -, various cortical areas (Fig. *6A-H*, *left*), and the amygdala (Fig. 6*F*, *left*). On the other hand, mPFC_VTA_ innervation was strongest in the lateral (LS) and medial septum (MS; Fig. 6*C*, *right*), the hypothalamus (HT), the bed nucleus of the stria terminalis (BNST; Fig. 6*D-F*, *right*), the midline thalamic nuclei (Fig. 6*E-F*), the zona incerta (ZI; Fig. 6*F*, *right*) and various tectal (Fig. 6*G-H*, *right*), tegmental – including the VTA – (Fig. 6*G-I*, *right*) and pontine regions (Fig. 6*G-I*, *right*). Taken together, our investigation revealed that mPFC_NAc_ and mPFC_VTA_ populations differ in their projection patterns not only in the NAc and VTA, but throughout the brain.

**Figure 6.**
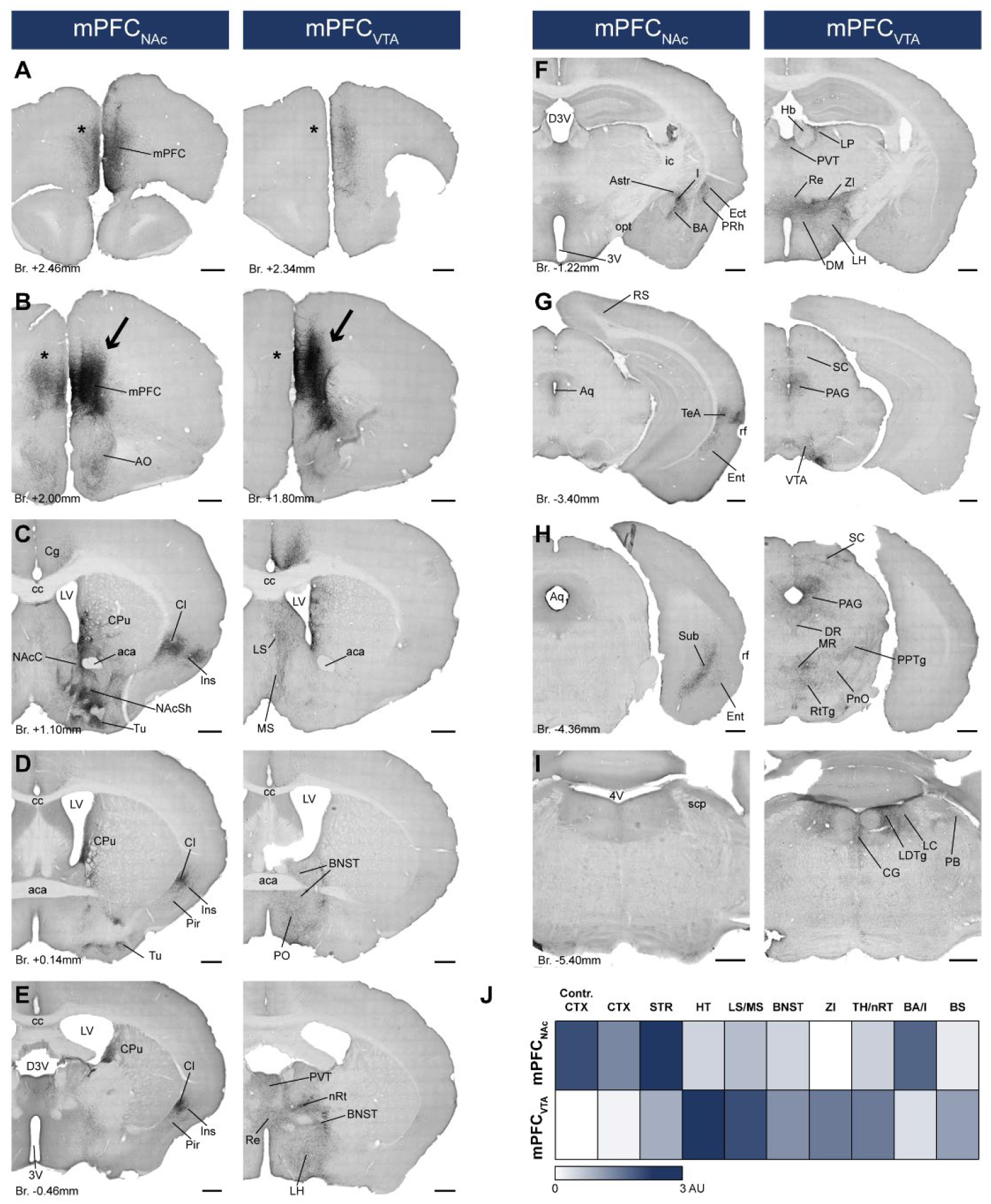
mPFC_NAc_ and mPFC_VTA_ neurons possess different efferent connections. **A-I,** Brightfield images showing the distribution AAV-DIO-mCherry labelled axons visualized with IHC_DAB-Ni_ at different AP levels. Arrows indicate AAV injection sites in the mPFC (**B**). Note the clear difference between the mPFC_NAc_ (left column) and mPFC_VTA_ (right column) populations, most prominently in the striatum (**C**), different cortical areas (**A-H**), the hypothalamus (**D-F**) and the brainstem (**F-I**) – including the VTA (**G**). Note the almost complete lack of contralateral cortical projection in mPFC_VTA_ animals as opposed to mPFC_NAc_ animals (**A-B**, asterisks). For experimental design see Fig. 6A. **J,** Summary table showing the innervation intensities of mPFC_NAc_ (top row) and mPFC_VTA_ (bottom row) populations in different brain regions. Darker color indicates stronger innervation. Scale bars: 500 μm. 3V, 3rd ventricle; 4V, 4th ventricle; aca, anterior commissure, anterior part; AO, anterior olfactory nucleus; Astr, amygdalostriatal transition area; Aq, aqueduct; BA, basolateral amygdaloid nucleus; BNST, bed nucleus of the stria terminalis; BS, brainstem; cc, corpus callosum; CG, central gray; CPu, caudate putamen; CTX, cortex; D3V, dorsal 3rd ventricle; DM, dorsomedial hypothalamic nucleus; DR, dorsal raphe; Ect, ectorhinal cortex; Ent, entorhinal cortex; Hb, habenula; I, intercalated amygdalar nuclei; ic, internal capsule; Ins, insular cortex; LH, lateral hypothalamus; LC, locus coeruleus; LDTg, laterodorsal tegmental nucleus; LP, lateral posterior thalamic nucleus; LS, lateral septum; LV, lateral ventricle; MR, medial raphe; MS, medial septum; nRT, reticular thalamic nucleus; PVT, paraventricular thalamic nucleus; VDB, nucleus of the vertical limb of the diagonal band; VP, ventral pallidum; opt, optic tract; PAG, periaqueductal gray; PB, parabrachial nucleus; Pir, piriform cortex; PnO, pontine reticular nucleus, oral part; PO, preoptic area; PPTg, pedunculopontine tegmental nucleus; PRh, perirhinal cortex; Re, reuniens thalamic nucleus; rf, rhinal fissure; RS, retrosplenial cortex; RtTg, reticulotegmental nucleus of the pons; SC, superior colliculus; scp, superior cerebellar peduncle; STR, striatum; Sub, subiculum; TeA, termporal association cortex; TH, thalamus; Tu, olfactory tubercule; ZI, zona incerta.

**Figure 7.**
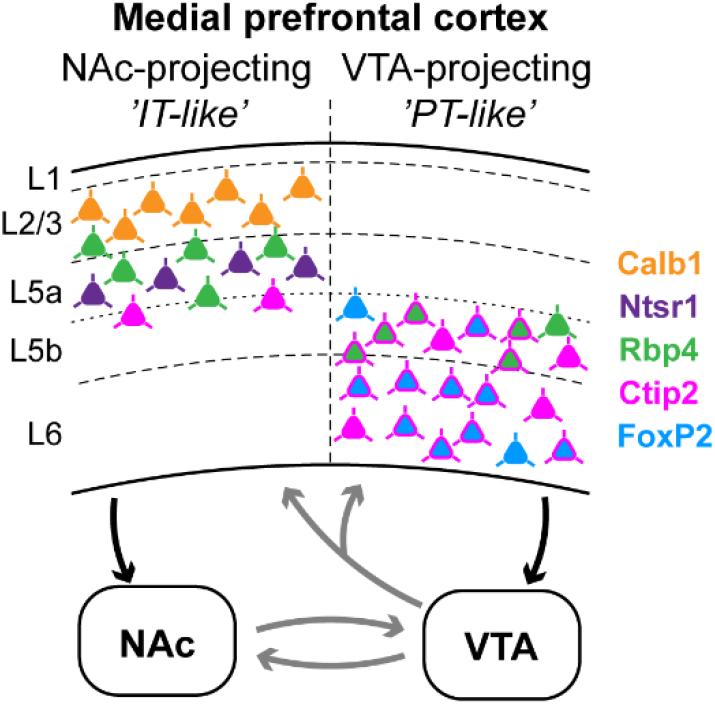
Summary: molecular characteristics and laminar distribution of the two identified projection groups in the mPFC. Neurons that innervate the NAc (‘IT-like’) are mostly localized in the upper layers of the mPFC (L2/3-5a) and express Calb1 (green), Ntsr1 (purple), Rbp4 (orange) and, to a lesser extent, Ctip2 (magenta). mPFC cells that innervate the VTA (‘PT-like’) are mostly localized in the deeper layers (L5b-6) and express Ctip2, FoxP2 (cyan) and Rbp4. Connections between NAc and VTA, and ascending VTA pathways (gray arrows) are based on literature data (see Introduction, Discussion).

**Table 5.**
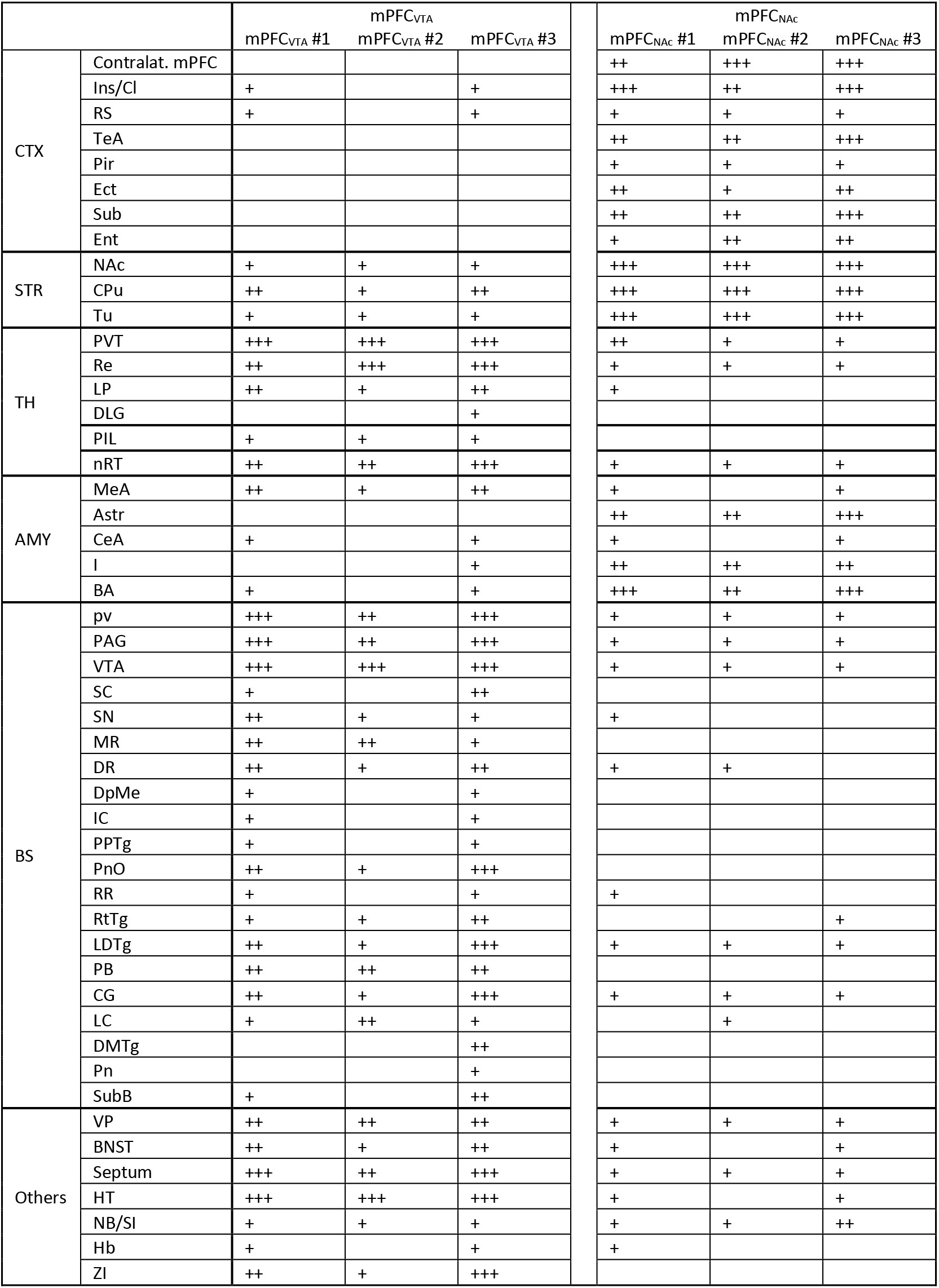
Whole-brain mapping data showing the axon-densities in different brain regions in the mPFC_NAc_ and mPFC_VTA_ animals (n = 3-3 mice). AMY, amygdala; Astr, amygdalostriatal transition area; BA, basolateral amygdaloid nucleus; BNST, bed nucleus of the stria terminalis; BS, brainstem; CeA, central amygdaloid nucleus; CG, central gray; Cl, claustrum; CPu, caudate putamen; CTX, cortex; DLG, dorsal lateral geniculate nucleus; DMTg, dorsomedial tegmental area; DpMe, deep mesencephalic nucleus; DR, dorsal raphe; Ect, ectorhinal cortex; Ent, entorhinal cortex; Hb, habenula; HT, hypothalamus; I, intercalated amygdalar nuclei; IC, inferior colliculus; Ins, insular cortex; LC, locus coeruleus; LDTg, laterodorsal tegmental nucleus; LP, lateral posterior thalamic nucleus; MeA, medial amygdaloid nucleus; MR, medial raphe; NB, basal nucleus; nRT, reticular thalamic nucleus; PVT, paraventricular thalamic nucleus; VP, ventral pallidum; PAG, periaqueductal gray; PB, parabrachial nucleus; PIL, posterior intralaminar thalamic nucleus; Pir, piriform cortex; Pn, pontine nuclei; PnO, pontine reticular nucleus, oral part; PPTg, pedunculopontine tegmental nucleus; pv, periventricular fiber system; Re, reuniens thalamic nucleus; RR, retrorubral nucleus; RS, retrosplenial cortex; RtTg, reticulotegmental nucleus of the pons; SC, superior colliculus; SI, substantia innominata; STR, striatum; Sub, subiculum; SubB, subbrachial nucleus; TeA, termporal association cortex; TH, thalamus; Tu, olfactory tubercule; ZI, zona incerta.

|  |  | mPFC<sub>VTA</sub> |  |  | mPFC<sub>NAC</sub> |  |  |
| --- | --- | --- | --- | --- | --- | --- | --- |
|  |  | mPFC<sub>VTA</sub> #1 | mPFC<sub>VTA</sub> #2 | mPFC<sub>VTA</sub> #3 | mPFC<sub>NAC</sub> #1 | mPFC<sub>NAC</sub> #2 | mPFC<sub>NAC</sub> #3 |
| CTX | Contralat. mPFC |  |  |  | ++ | +++ | +++ |
|  | Ins/Cl | + |  | + | +++ | ++ | +++ |
|  | RS | + |  | + | + | + | + |
|  | TeA |  |  |  | ++ | ++ | +++ |
|  | Pir |  |  |  | + | + | + |
|  | Ect |  |  |  | ++ | + | ++ |
|  | Sub |  |  |  | ++ | ++ | +++ |
|  | Ent |  |  |  | + | ++ | ++ |
| STR | NAC | + | + | + | +++ | +++ | +++ |
|  | CPu | ++ | + | ++ | +++ | +++ | +++ |
|  | Tu | + | + | + | +++ | +++ | +++ |
| TH | PVT | +++ | +++ | +++ | ++ | + | + |
|  | Re | ++ | +++ | +++ | + | + | + |
|  | LP | ++ | + | ++ | + |  |  |
|  | DLG |  |  | + |  |  |  |
|  | PIL | + | + | + |  |  |  |
|  | nRT | ++ | ++ | +++ | + | + | + |
| AMY | MeA | ++ | + | ++ | + |  | + |
|  | Astr |  |  |  | ++ | ++ | +++ |
|  | CeA | + |  | + | + |  | + |
|  | I |  |  | + | ++ | ++ | ++ |
|  | BA | + |  | + | +++ | ++ | +++ |
| BS | pv | +++ | ++ | +++ | + | + | + |
|  | PAG | +++ | ++ | +++ | + | + | + |
|  | VTA | +++ | +++ | +++ | + | + | + |
|  | SC | + |  | ++ |  |  |  |
|  | SN | ++ | + | + | + |  |  |
|  | MR | ++ | ++ | + |  |  |  |
|  | DR | ++ | + | ++ | + | + |  |
|  | DpMe | + |  | + |  |  |  |
|  | IC | + |  | + |  |  |  |
|  | PPTg | + |  | + |  |  |  |
|  | PnO | ++ | + | +++ |  |  |  |
|  | RR | + |  | + | + |  |  |
|  | RtTg | + | + | ++ |  |  | + |
|  | LDTg | ++ | + | +++ | + | + | + |
|  | PB | ++ | ++ | ++ |  |  |  |
|  | CG | ++ | + | +++ | + | + | + |
|  | LC | + | ++ | + |  | + |  |
|  | DMTg |  |  | ++ |  |  |  |
|  | Pn |  |  | + |  |  |  |
|  | SubB | + |  | ++ |  |  |  |
| Others | VP | ++ | ++ | ++ | + | + | + |
|  | BNST | ++ | + | ++ | + |  | + |
|  | Septum | +++ | ++ | +++ | + | + | + |
|  | HT | +++ | +++ | +++ | + |  | + |
|  | NB/SI | + | + | + | + | + | ++ |
|  | Hb | + |  | + | + |  |  |
|  | ZI | ++ | + | +++ |  |  |  |

## Discussion

Here we described the molecular, neurochemical, and anatomical characteristics of mPFC neurons innervating the mesocorticolimbic system. We found that most mPFC neurons projecting to the NAc and the VTA were localized in the same subregions, but in different layers. Accordingly, mPFC_NAc_ and mPFC_VTA_ neuron populations showed minimal overlap at the cellular level, expressed different combination of layer-specific molecular markers and their efferent connections showed clear differences throughout the brain. While mPFC_NAc_ neurons mostly innervated ipsi- and contralateral cortical, striatal and amygdalar regions, mPFC_VTA_ axons were most abundant in various ipsilateral diencephalic and mesencephalic areas.

Generally, mPFC_NAc_ and mPFC_VTA_ neurons were found in the same subregions, namely the PL, IL, MO, Cg1, DP and DTT, confirming previous results (Gabbott et al., 2005). However, one notable difference emerged between the two populations. While mPFC_NAc_ neurons formed one, mostly continuous cluster with the highest number of cells in the PL, IL, and MO, mPFC_VTA_ neurons formed two visually distinct laminar clusters: one in the middle and another in the deeper part of the mPFC.

Regarding their laminar distribution, mPFC_NAc_ neurons were mostly found in the superficial layers (L2/3 and L5a), as previously reported (Kim et al., 2017). Traditionally, most striatum-projecting cortical neurons belong to the IT projection group (Harris and Shepherd, 2015). High ratio of Calb1 expressing neurons in the L2/3 (^~^70%) and strong innervation of the NAc in the Calb1-Cre animals also suggest their IT-like nature, since Calb1 is considered to be an IT-marker (Harris et al., 2019). The functional importance of these L2/3 mPFC cells has been shown by Shrestha and colleagues (2015) demonstrating that their genetic perturbation leads to augmented depressive behavior in response to stressful events, possibly via the NAc-hypothalamic pathway.

In addition to Calb1, Rbp4 - a genetic marker for both IT and PT neurons (Rojas-Piloni et al., 2017; Harris et al., 2019) -, was also expressed to some extent in the L2/3 besides the L5 of the mPFC. Accordingly, these cells provided strong input to NAc. Surprisingly, despite their relatively low number, mPFC neurons expressing Ntsr1, distributed only in the L5a, also heavily innervated the NAc. These observations indicate regional differences in the distribution of the Rbp4 and Ntsr1-expressing cortical neurons, since Rbp4 is known to be present in the L5, while Ntsr1 is a generally used marker for L6 CT neurons in other, mostly primary cortical regions (Jeong et al., 2016; Sundberg et al., 2018; Matho et al., 2021). We confirmed these results using the same viral tracing experimental approach and the same animal strains targeting the neighboring primary motor cortex to exclude the possibility of a faulty mouse/viral strain. In fact, Rbp4 cells were exclusively localized in the L5 of M1. Furthermore, Ntsr1 neurons were only distributed in the L6 of the M1 and innervated the thalamus but not the striatum (data not shown). These results indicate that some molecular markers have distinct laminar distribution and projection patterns in primary and higher order cortical areas.

Further supporting this notion, we demonstrated that Ctip2, which is generally present in PT neurons of the L5b-L6 (Arlotta et al., 2005; Ueta et al., 2014; Kim et al., 2017) was expressed in about one-fifth of all mPFC_NAc_ (IT-like) neurons. This suggests that either some PT-like mPFC neurons innervate the striatum or, alternatively, some IT-like neurons express Ctip2 in the mPFC. Previous results reported that PT neurons in the motor cortex can weakly innervate the striatum (Economo et al., 2018; Matho et al., 2021) supporting the first option. However, to the best of our knowledge, there is no direct evidence for the complete absence of Ctip2-expression in IT neurons, so we cannot completely rule out the second possibility either.

While mPFC_NAc_ neurons were present rather superficially, mPFC_VTA_ neurons were mostly localized in the deeper layers, namely in L5b and L6 (Geisler and Zahm, 2005) and the vast majority (^~^95%) of them expressed Ctip2. Furthermore, Rbp4 neurons – shown to have reinforcing effect (Pan et al., 2021) – innervated the VTA and the NAc with a similar intensity. If we assume that IT- and PT-like Rbp4 neurons are spatially separated (in L2/3-L5a and L5b, respectively), and that IT-like neurons innervate the NAc but not the VTA, then, these results suggest that mPFC_VTA_ neurons have a PT-like phenotype. However, FoxP2, a L6 CT neuron marker (Kast et al., 2019; Matho et al., 2021) was also expressed by almost half of all mPFC_VTA_ cells. This observation was confirmed by cell-specific viral tracing in the FoxP2-Cre mouse strain, where labelled neurons were found in the L6 and projected heavily to the VTA and to the thalamus (data not shown), resembling a mixed PT-CT population. Accordingly, axons of the CAV2-Cre-labelled mPFC_VTA_ neurons collateralized to the thalamus as well. In contrast, FoxP2 neurons in the M1 cortex showed clear CT phenotype (data not shown), as it was previously reported (Matho et al., 2021). These results strengthened our previous assumption that some cell-types have different anatomical phenotype in primary and prefrontal cortical regions.

The different laminar distribution and molecular characteristics of mPFC_NAc_ and mPFC_VTA_ neurons suggest that these populations are mostly separated. However, previous publications yielded contradictory results about the target-selectivity of mPFC neurons, which can be resolved, if we consider that multiple projection was found to be high when the experiments were carried out in one neuron population (e.g., only IT or only PT neurons) (Thierry et al., 1983; Ferino et al., 1987; Cassell et al., 1989; Vázquez-Borsetti et al., 2011; Rojas-Piloni et al., 2017), but low when the experiments involved mixed populations (e.g., PT and IT neurons) (Pinto and Sesack, 2000; Gabbott et al., 2005; Morishima and Kawaguchi, 2006; Kim et al., 2017; Otis et al., 2017; Cruz et al., 2021). Gao et al. (2020) found relatively high overlap between NAc- (IT) and VTA (PT) projecting neurons in the Cg1, however, examining only a relatively small sample size. Recent publications (Callaway et al., 2021; Muñoz-Castañeda et al., 2021; Peng et al., 2021) showing extensive collateralization of fully reconstructed IT and PT neurons also suggest that cortical pyramidal neurons have a multiple-projecting nature. Collectively, these results imply that a single mPFC neuron is not target-selective, but target-preference can be identified at class, or subclass level. We directly tested this and found that retrogradely labelled mPFC_Nac_ and mPFC_VTA_ neurons showed minimal overlap (<2%), indeed. Furthermore, using CAV2-Cre-mediated viral tracing we demonstrated that mPFC_NAc_ cells innervate the NAc approximately ten times stronger than the VTA. On the other hand, mPFC_VTA_ neurons also showed clear preference (3.5-fold) for the VTA over the NAc. Considering that mPFC innervates the VTA with a relatively sparse axon-arborization (Carr and Sesack, 2000; Geisler and Zahm, 2005), these result further support that these populations are rather non-overlapping at the single-cell level. Complete projection pattern analysis revealed that these populations tend to innervate different areas throughout the brain. Specifically, mPFC_NAc_ neurons showed IT-like projection pattern (mainly ipsi- and contralateral cortical, amygdalar and striatal targets), while mPFC_VTA_ efferents resembled PT neurons (mainly ipsilateral mesencephalic and diencephalic targets).

In general, mPFC_NAc_ neurons participate in a range of reward-related tasks. For example, activation of mPFC_NAc_ neurons suppresses reward seeking in a conflicting situation (Kim et al., 2017). On the other hand, others reported that optical stimulation of mPFC_NAc_ neurons promote conditioned reward seeking (Otis et al., 2017). In accordance, Britt et al. (2012) demonstrated that optical stimulation of mPFC terminals in the NAc can facilitate self-stimulation, although Stuber and colleagues (2011) reported the lack of such effect. Therefore, it seems plausible that there is a topographical segregation within the mPFC-to-NAc pathway with different functional properties or different cell-types conveying different behavioral information, or the combination of both. Similarly, it was previously reported that mPFC neurons can excite and inhibit VTA dopamine neurons equally (Lodge, 2011), which also suggests functional separation within the mesocorticolimbic system. Recent findings of topographically biased input-output connectivity of different mPFC (Cruz et al., 2021) and VTA dopamine neurons (Aransay et al., 2015; Beier et al., 2015), as well as high topographic precision in corticostriatal pathways (Hooks et al., 2018) further support this suggestion. So, cell-specific studies are needed to completely clarify the functional complexity of these pathways.

Taken together, mPFC_NAc_ and mPFC_VTA_ populations are rather non-overlapping and their afferent connectivity shows IT- and PT-like features, respectively. However, high CT marker (FoxP2) expression in mPFC_VTA_ neurons, as well as PT (Ctip2) and CT (Ntsr1) marker expression in mPFC_NAc_ neurons indicate that the traditional IT-PT-CT classification is less evident in the mPFC. In accordance, recent publications also demonstrated high genetic diversity of cortical neurons (Bakken et al., 2021; Callaway et al., 2021; Matho et al., 2021; Zhang et al., 2021), even within projection neuron classes in primary cortices. Therefore, in the future, understanding the versatility of prefrontal cortical influence over mesocorticolimbic functions requires a combination of molecular-, cellular-, laminar- and region-specific approaches.

### Anatomical considerations

It is generally accepted that the rodent mPFC is anatomically homologous to the primate anterior cingulate cortex (Russo and Nestler, 2013; Vogt and Paxinos, 2014). However, there are notable nomenclatural inconsistencies (Laubach et al., 2018; Le Merre et al., 2021) in the rodent mPFC literature (Lodge, 2011; Bossert et al., 2012; Adhikari et al., 2015; Shrestha et al., 2015; Warren et al., 2019; Lichtenberg et al., 2021). For example, the exact definition of the PrL subregions greatly varies between publications, just like the distinction between dorsal and ventral mPFC. Such inaccuracies can contribute to the still abundant contradictions in the literature and complicate the proper interpretation of the results.

To overcome these setbacks, we combined multiple IHC_Fluo_ against different molecular markers that can (1) delineate the borders between different subregions (PV, Calb1) (van Brederode et al., 1991; Sun et al., 2002; Akhter et al., 2014; Mátyás et al., 2014) and (2) clearly define cortical layers in the mPFC (Calb1, Ctip2, FoxP2) (Ferland et al., 2003; Kim et al., 2017). We always used these markers to locate injection sites and labelled neurons within the mPFC. Reliable primary antibodies raised in several different species against all of these markers are commercially available and they can be combined easily. Therefore, we suggest the general adoption of this method to precisely define and separate mPFC subregions and layers in future studies.

## Acknowledgements

The authors thank Tamás Herczeg, Réka Erdős, Anna Fehér and Dóra Zsíros for laboratory assistance, the Institute of Enzymology of the Research Centre for Natural Sciences for the confocal microscope, Dóra Zelena for the CAV2-Cre virus, Norbert Hájos for providing antibodies and László Acsády, Péter Barthó and Balázs Rózsa for the Cre animals. We also thank Norbert Hájos for comments and discussions about the manuscript. This work was supported by the Ministry of Innovation and Technology of Hungary from the National Research, Development and Innovation Fund (FK124434, K138836 and KKP126998 to F.M), by the Hungarian Brain Research Program (2017-1.2.1-NKP-2017-00002 to F.M.) by the New National Excellence Program of the Ministry for Innovation and Technology (ÚNKP-21-5-ÁTE-2 to F.M.; ÚNKP-18-3-II-BME-55, ÚNKP-20-3-II-BME-24 and ÚNKP-21-3-II-BME-61 to Á.B.). F.M. is a János Bolyai Research Fellow.

## Materials and Methods

### Animals

Adult (3-5 months old, male and female) wild-type, Rbp4-Cre (Tg(Rbp4-cre)KL100Gsat, RRID: MMRRC_037128-UCD, gift from L. Acsády), Thy1-Cre (FVB/N-Tg(Thy1-cre)1Vln/J, RRID: IMSR_JAX:006143; gift from B. Rózsa), Calb1-Cre (B6;129S-Calb1^tm2.1(cre)Hze^/J, RRID: IMSR_JAX:028532), Ntsr1-Cre (Tg(Ntsr1-cre)GN220Gsat, RRID: MMRRC_017266-UCD a gift from P. Barthó) and FoxP2-Cre mice (B6.Cg-Foxp2^tm1.1(cre)Rpa^/J, RRID: IMSR_JAX:030541) were used for the experiments. Animals were group housed in a humidity- and temperature-controlled environment. Animals were entrained to a 12 h light/dark cycle (light phase from 07:00 AM) with food and water available *ad libitum*. All procedures were approved by the Regional and Institutional Committee of the Research Centre for Natural Sciences.

### Stereotactic surgeries

#### Classical retrograde tracing

All animals were anesthetized under ketamine-xylazine (5:1, 3x dilution, ketamine: 100 mg/kg; xylazine: 4 mg/kg) during all anatomical surgeries. Single and double retrograde tracing surgeries were carried out with 0.5% Cholera Toxin B subunit (CTB; List Biological Laboratories: 104) and/or 2% Fluoro-Gold (FG; Fluorochrome LLC) to reveal the prefrontal cortical source of NAc (AP/L/DV: +1.4/±0.8/3.9-4.2) and VTA (AP/L/DV: −3.3/±0.3/4.0-4.2) innervation. Tracers were iontophoretically injected (7-7 sec on/off duty cycle, 3-5 μA, for 5-10 min) with IonFlow Bipolar electrophoretic equipment (Supertech Instruments Hungary). After all surgeries, animals received Rimadyl (Carprofen, 1.4 mg/kg).

For anatomical analysis, after 7 days of survival time, mice were perfused transcardially first with saline (^~^50 ml), then, with ^~^150 ml of fixative solution containing 4% paraformaldehyde (Sigma) in 0.1 M phosphate buffer (PB). Animals in which the injections sites or tracer tracks reached regions that could affect labelling (e.g.: caudate putamen, substantia nigra, ventral pallidum) were excluded from further analysis.

#### Identification of different brain regions and cortical layers

We used different neurochemical markers to identify brain regions of interest and to separate cortical layers in the tissue samples labelled with fluorescent immunohistochemistry (IHC_Fluo_). Calbindin (Calb1) staining (see below) was used to delineate the core (strong Calb1 expression) and shell (weak Calb1 expression) region of the NAc (Jongen-Rêlo et al., 1994), and tyrosine hydroxylase (TH) staining for the VTA (Oades and Halliday, 1987; Morales and Margolis, 2017). Layer 2/3 (L2/3) of the cerebral cortex was identified using Calb1 staining (van Brederode et al., 1991; Sun et al., 2002), while L6 with forkhead box protein P2 (FoxP2) staining (Ferland et al., 2003). COUP-TF-interacting protein 2 (Ctip2) staining was used to label L5b and L6 (DeNardo et al., 2015) (Fig. 1-1).

We used the 2nd Edition of the Mouse Brain is Stereotaxic Coordinates by Paxinos and Franklin (2001) as a reference, because the vast majority of mPFC literature uses this nomenclature. In comparison with the newest, 5th edition (Franklin and Paxinos, 2019), the mPFC region we defined as prelimbic cortex (PrL) is approximately equivalent to the A32 area, the infralimbic cortex (IL) to the A25, and the rostral aspects of the cingulate cortex, area 1 and 2 (Cg1-2) to the A24b and A24a, respectively. The secondary motor (M2), medial orbital (MO), dorsal peduncular cortex (DP) and dorsal tenia tecta (DTT) regions have not changed significantly between the two editions.

#### Anterograde viral tracing

For cell type-specific anterograde viral tracing AAV5.EF1a.DIO.eYFP.WPRE.hGH (30-100 nl; Penn Vector Core; #27056-AAV5; titer: 5*10^12^ GC/ml) or AAV5-EF1a-DIO-mCherry viruses (30-100 nl; UNC Vector Core; #50462; titer: 7*10^12^ GC/ml) were injected at a rate of 0.5-1 nl/sec into mPFC (AP/L/DV: +1.7-1.9/±0.3/2.1-1.6 mm) and M1 (AP/L/DV: +1.4/±1.6/1.3-1.0 mm) using a Nanoliter Injector (World Precision Instruments, FL, USA).

Animals were perfused (see above) after 4-6 weeks of survival time. Viral expression was always analyzed after IHC_Fluo_-enhancement (Fig. 3-1) (Falcy et al., 2020), even for eYFP (see below).

#### Intersectional retro-anterograde viral tracing

In order to selectively label NAc-projecting (mPFC_NAc_) and VTA-projecting mPFC cells (mPFC_VTA_), we injected Canine adenovirus type 2 carrying Cre-recombinase gene (CAV2-Cre, CMV promoter, titer: 2.5*10^10^ pp/ml, Plateforme de Vectorologie de Montpellier, France; a gift from D. Zelena) into the NAc (*n* = 3 animals) or VTA (*n* = 3 animals) (see coordinates above) of wild-type animals, mixed with 5% biotinylated dextrane amine (BDA, MW: 10.000, Molecular Probes: D1956, RRID: AB_2307337; 1:1; 80-120 nl/animal; 1nl/sec). Note that BDA was used to locate the tip of the injecting pipette (Fig. 5B1, C1), not the whole extent of viral diffusion. At the same time, the mPFC (see coordinates above) of the same animals was injected with AAV5-EF1a-DIO-mCherry (see details above). After 6 weeks of survival, animals were perfused, and their brains were processed for further analysis (see above).

### Tissue processing and immunohistochemistry

Tissue blocks were cut on a VT1200S Vibratome (Leica) into 50 μm coronal sections. Free-floating sections were intensively washed with 0.1 M PB. All antibodies were diluted in 0.1 M PB. For fluorescent labeling, sections were first treated with a blocking solution containing 10% normal donkey serum (NDS, Sigma-Aldrich: S30-M) or 10% normal goat serum (NGS, Vector: S-1000, RRID: AB_2336615) and 0.5% Triton-X in 0.1 M PB for 30 minutes at room temperature (RT).

#### Fluorescent immunohistochemistry

Sections were incubated in primary antibody solution overnight at RT or for 2-3 days at 4°C. The following primary antibodies were used: green fluorescent protein (GFP, chicken, Life Technology: A10262, RRID: AB_2534023; 1:2000), mCherry (mCherry; rabbit, BioVision: 5993-100, RRID: AB_1975001; 1:2000), red fluorescent protein (RFP; rat, Chromotek: 5F8, RRID: AB_2336064; 1:2000) FoxP2 (mouse, Merck Millipore: MABE415, RRID: AB_2721039; 1:2000; Invitrogen: MA5-31419, RRID: AB_2787055; 1:2000; rabbit, Abcam: ab16046, RRID: AB_2107107; 1:500), Calb1 (rabbit, SWANT: CB38, RRID: AB_10000340; 1:2000; mouse, SWANT: 300, RRID: AB_10000347; 1:2000; chicken, Synaptic Systems: 214 006, RRID: AB_2619903; 1:2000), TH (mouse, Immunostar: 22941, RRID: AB_572268; 1:8000), FG (rabbit, FluoroChrome, 1:50.000; guinea pig, Protos Biotech: NM-101, RRID: AB_2314409; 1:5000), CTB (goat, List Biological Laboratories: 703; 1:20.000), parvalbumin (PV; mouse, SWANT: PV 235, RRID: AB_10000343; 1:2000) and Ctip2 (rat, Abcam: ab18465, RRID: AB_2064130; 1:500).

For IHC_Fluo_ staining, after primary antibody incubation, sections were treated with the following secondary IgGs (1:500; 2 hours at RT): Alexa 488-conjugated donkey anti-rabbit (DAR-A488; Jackson: 711-545-152, RRID: AB_2313584), donkey anti-mouse (Jackson: 715-545-150, RRID: AB_2340846), goat anti-chicken (Molecular Probes: A11039, RRID: AB_142924), donkey anti-guinea pig (Jackson: 706-545-148, RRID: AB_2340472); Alexa 555-conjugated donkey anti-goat (Molecular Probes: A21432, RRID: AB_141788), donkey anti-mouse (Molecular Probes: A31570, RRID: AB_2536180), donkey anti-rat (Southern Biotech: 6430-32, RRID: AB_2796359); Cy3-conjugated donkey anti-rabbit (Jackson: 715-165-152, RRID: AB_2307443), donkey anti-mouse (Jackson: 715-165-151, RRID: AB_2340813); Alexa 594-conjugated donkey anti-mouse (Molecular Probes: A21203, RRID: AB_141633), donkey anti-rabbit (Molecular Probes: A21207, RRID: AB_141637), Alexa 647-conjugated donkey anti-mouse (Jackson: 715-605-151, RRID: AB_2340863; Invitrogen: A-31571, RRID: AB_162542) or donkey anti-rabbit (Jackson: 711-605-152, RRID: AB_2492288).

When necessary, staining was enhanced after primary antibody incubation with biotinylated secondary antibodies (biotinylated horse anti-goat IgG, Vector Laboratories: BA-9500, RRID: AB_2336123; 1:300; biotinylated goat anti-rabbit – bGAR, Vector Laboratories: BA-1000, RRID: AB_2313606; 1:300; biotinylated goat anti-guinea pig, Vector Laboratories: BA-7000, RRID: AB_2336132; 1:300; 1.5h, RT), Elite Avidin-Biotin Complex (eABC, 1:300, Vector Laboratories: PK-6100, RRID: AB_2336819; 1.5h, RT) and streptavidin conjugated fluorescent antibodies (SA-A488, Jackson: 016-540-084, RRID: AB_2337249; 1:2000; SA-Cy3, Jackson: 016-160-084, RRID: AB_2337244; 1:2000; SA-A647, Jackson: 016-600-084, RRID: AB_2341101; 1:2000; 2 hrs, RT). All fluorescent slices were mounted in Vectashield (Vector Laboratories: H-1000, RRID: AB_2336789). To reveal the CAV2-Cre/BDA injection site we used eABC (see above) and SA-A488 or SA-A647 (see above).

#### Immunoperoxidase staining

For the whole-brain projection pattern analysis of the CAV2-Cre animals, we also performed immunoperoxidase staining and used nickel-amplified diaminobenzidine (DAB) technique (DAB-Ni; IHC_DAB-Ni_). Every 6^th^ section (thus, at 300 μm resolution, from Br. +3.10 to −8.00 mm) was treated first with 1% H2O2 solution for 10 minutes, then, after intensive washing, in 10% NDS and 0.2% Triton-X solution as a blocking serum (30 mins, RT). After primary antibody incubation (mCherry, see above), slices were incubated in biotinylated secondary antibody (bGAR) and eABC (see above). Then we developed DAB-Ni for 5 minutes. Sections were then dehydrated in xylol (2*10 mins) and mounted in DePex (Serva, Heidelberg, Germany).

#### Viral signal amplification

To compare native mCherry expression to IHC_Fluo_ and IHC_DAB-Ni_ enhancement, we stained slices from the CAV2-Cre experiments with primary antibody against mCherry and DAR-A488 (see above) (Fig. 3-1*A-B*). Then we captured confocal images (see below) from the same brain regions in two channels (i.e., A488 and mCherry). For better visualization, we recolorized the A488 channel at Fig. 3-1*A2*, *B2*. Next, we stained the neighboring slices (i.e., 50 μm apart) with IHC_DAB-Ni_ against mCherry (see above) and captured them with brightfield microscopy (see below) (Fig. 3-1*C-D*).

### Microscopy

Fluorescent sections were first analyzed with epifluorescent microscope (Leica DM 2500, Leica Microsystems GmbH; Camera: Olympus DP73, CellSens Entry 1.16, Olympus Corporation) with low magnification (2.5x N PLAN 2.5x/0.07 ∞/-/OFN25, 5x HCX FL PLAN 5x/0.12 ∞/-/B) to find injection sites and labelled cells. Higher magnification (10x Plan Apochromat 10x/0.45 M27; 20x Plan Apochromat 20x/0.8 M27, 63x Plan Apochromat 63x/1.4 Oil DIC M27) images were taken with confocal microscope (Zeiss LSM 710; Zeiss ZEN 2010B SP1 Release version 6.0; Carl Zeiss Microimaging GmbH). Brightfield imaging and whole brain projection analysis was completed with a PANORAMIC MIDI II (20x (NA 0.8); 3DHistech, Hungary) device and the manufacturer’s official software (CaseViewer 2.4) for every 6th slice (i.e., at 300 μm resolution).

#### Colocalization

In order to reveal the proportion of FoxP2-, Ctip2- and Calb1-positive cells among retrogradely (FG/CTB) labelled mPFC_NAc_ and mPFC_VTA_ cells, we captured 20x magnification confocal Z-stack (step size: 5 μm) imaging of double-labelled fluorescent sections (3-4 slices/animal, *n* = 3-3 animals). Labelled cells were then manually analyzed with ImageJ (NIH). Only cells visible in two separate sections with a visible nucleus were analyzed. The same protocol was used to identify double-labelled cells in the double retrograde tracing experiments (*n* = 3 animals).

#### Axon density analysis

We sought to compare mPFC axon densities in the NAc and VTA in the CAV2-Cre injected mPFC_NAc_, mPFC_VTA_, and AAV5-EF1a-DIO-mCherry injected Rbp4- (mPFC_Rbp4_) and Thy1-Cre (mPFC_Thy1_) samples using high-magnification (63x) confocal Z-stacks (step size: 0.27 μm). In the VTA we captured three stacks in each animal (*n* = 3 in each strain) at three different AP levels between Bregma −3.10 and −3.80 mm. In the NAc, we captured five-five stacks in the same animals as for the VTA at three different AP levels between Bregma +1.00-1.80 mm. We aimed to capture stacks where axon density was visibly the highest at each AP levels in each region.

We analyzed the confocal stacks using a custom made automatic ImageJ macro (Mátyás et al., 2018). The macro calculated the axon length for each stack and the total axon length was summated for each brain region in each animal. Then, the total axon length was compared to the summated stack volume (ROI area * number of slices * step size = total volume) for each brain region to calculate the relative axon density (RAD = total axon length/total volume). Then, the ratio of RAD_VTA_/RAD_NAc_ (RAD_(VTA/NAc)_) was calculated for each animal, where RAD_(VTA/NAc)_ = 1 means that the two areas are equally innervated.

### Statistical analysis

Values are given as mean ± SD in all figures. *n* represents number of animals; *N* represents cell counts. We used SPSS Statistics (ver. 27.0.1.0., IBM) to analyze the axon-density data. We used one-way ANOVA method with LSD post-hoc test to compare relative axon-density values after testing for the homogeneity of variances. *\*p* < 0.05, *\*\*\*p* < 0.001.

